# Matrix-rigidity cooperates with biochemical cues in M2 macrophage activation through increased nuclear deformation and chromatin accessibility

**DOI:** 10.1101/2024.02.13.579995

**Authors:** Seung Jae Shin, Buuvee Bayarkhangai, Khaliunsarnai Tsogtbaataar, Meng Yuxuan, Sang-Hyun Kim, Yong-Jae Kim, Daesan Kim, Dong-Hwee Kim, Jung Hwan Lee, Jeongeun Hyun, Hae-Won Kim

**Author notes:** Corresponding authors: DH Kim, JH Lee, J Hyun, and HW Kim.

## Abstract

Macrophages encounter a myriad of biochemical and mechanical stimuli across various tissues and pathological contexts. Notably, matrix rigidity has emerged as a pivotal regulator of macrophage activation through mechanotransduction. However, the precise mechanisms underlying the interplay between mechanical and biochemical cues within the nuclear milieu remain elusive. Here we elucidate how the increased matrix rigidity drives macrophages to amplify alternatively-activated (M2 phenotype) signalings cooperatively with biochemical cues (e.g., IL4/13) through altered nuclear mechanics. Notably, we found that reconstructed podosome-like F-actins and contractility induced nucleus deformation, opening nuclear pores, which facilitates nuclear translocation of the key transcription factor STAT6. Furthermore, the altered nuclear mechanics increased chromatin accessibility induced by H3K9 methylation, particularly of M2-associated gene promoters. These cooperative events of the mechano-chemical signaling at the cytoskeletal-to-nuclear domains facilitated M2 transcriptional activation and cellular functions. We further evidenced the rigidity-primed M2 macrophages were immunosuppressive and accumulated in stiffened tumor tissues. This study proposes a mechanism by which matrix mechanics crosstalks with biochemical signals to potentiate macrophage activation through nuclear mechanosensing and chromatin modifications, offering insights into macrophage mechanobiology and its therapeutic modulations.

## Introduction

Macrophages, as versatile immune cells, exhibit a broad range of phenotypes that encompass both pro-inflammatory and anti-inflammatory states in response to cues from the microenvironment ^1^. While macrophages display diverse phenotypes with high heterogeneity in various tissues ^2^, they are commonly categorized into classically activated (M1) and alternatively activated (M2) subtypes. M1 macrophages, activated by interferon-γ (IFN-γ) and lipopolysaccharide (LPS), engage in phagocytosis of bacteria and virus-infected cells and release pro-inflammatory cytokines to attract other immune cells ^3^. On the other hand, M2 macrophages, induced by interleukin-4 (IL4), IL10, and IL13, contribute to wound healing, tissue repair, and resolution of inflammation ^1,3,4^. The plasticity of macrophages is evident as they transition from M1 to M2 phenotypes to facilitate tissue repair following injury ^5^. This transition involves a decrease in the production of pro-inflammatory cytokines such as IL1β, IL6, and tumor necrosis factor-α (TNF-α) while elevating the expression of IL10, vascular endothelial growth factor (VEGF), and transforming growth factor-β1 (TGF-β1).

Macrophages are influenced not only by biochemical signals but also by altered mechanical cues within the body. Tissue-resident macrophages experience a diverse range of matrix rigidity in healthy and pathophysiological conditions ^6,7^. Additionally, monocyte-derived macrophages encounter mechanical stress during their migration from blood vessels to different tissues for replenishing resident macrophages and responding to inflammation ^2,8,9^. Existing studies reveal the mechanosensitivity of macrophages and the way how mechanical factors shape their functions. For instance, mouse bone marrow-derived macrophages (BMDMs) and human peripheral blood mononuclear cell-derived macrophages show enhanced adhesion and spreading on rigid substrates (hundreds of kPa) compared to compliant ones (∼1 to 10 kPa) ^10–12^. Research also indicates that macrophages cultured on rigid biomaterials potentiate to activate M1 responses upon IFN-γ and LPS treatment, explaining foreign body immune responses to implanted materials ^10,13,14^. Recent studies suggest that M2 macrophage activation can be enhanced on rigid matrices in the presence of IL4 and IL13 (IL4/13) ^11,14^, but the underlying mechanisms remain to be explored.

A comprehensive understanding of how the mechanical environment influences macrophage behavior could yield new strategies and targets for treating diseases, given that many diseases involve changes in mechanical properties of tissues. Tumors, for example, are characterized by increased collagen deposition and matrix rigidity, which contribute to disease progression and metastasis ^15^. Tumor-associated macrophages (TAMs), resembling M2 macrophages, are involved in this process ^15^. Matrix rigidity has been shown to enhance interactions between cancer cells and macrophages, expediting M2 macrophage activation in breast cancer microenvironments ^16^. A recent study by Kim, et al. ^11^ revealed that actin cytoskeleton polymerization and mechanotransduction, along with enhanced adhesion and spreading, drive M2 macrophage activation on rigid matrices by promoting the nuclear import of signal transducer and activator of transcription 6 (STAT6), a critical transcription factor (TF) for M2 activation. However, it remains to be investigated whether and how mechanical forces can affect chromatin organization and gene expression in order to regulate the phenotypic plasticity of macrophages in a timely and precise manner.

Hence, this study examines how chromatin organization changes with varying matrix rigidity during M2 macrophage activation, and how these changes interact with TFs to regulate gene expression. We find that macrophages cultured on more rigid matrices enhance M2 activation and anti-inflammatory phenotypes in the presence of IL4/13. Of note, more rigid conditions lead to less condensed chromatin, particularly within the nuclear lamina where genes are often silenced. More specifically, promoter and upstream regions of M2-associated genes, such as *Pparg*, *Tgm2*, and *Irf4*, become less condensed, while those of M1-associated genes, like *Il6* and *Tlr2*, become more condensed under higher rigidity conditions with IL4/13. Additionally, we demonstrate that M2 macrophages on rigid substrates form podosome-like actin structures crucial for nuclear deformation and mechanosensing. The nuclear deformation enables nuclear pore opening, facilitating nuclear import of phosphorylated STAT6 (pSTAT6), and chromatin rearrangement in M2-associated genomic loci. These findings underscore the roles of mechanical cue (matrix rigidity) synergized with biochemical cue (anti-inflammatory cytokines) in M2 macrophage activation, and the significance of nuclear mechanosensing for chromatin reorganization and enhanced nuclear transport of TF (pSTAT6), suggesting potential targets for disease treatment and tissue healing where macrophage activation is key involved, through the modulation of actin cytoskeleton-mediated nuclear mechanosensing.

## Results

### Matrix rigidity amplifies M2 macrophage activation cooperatively with biochemical cues

To explore the cooperative effect of mechanical and biochemical cues on M2 macrophage activation, we began with the characterization of primary mouse BMDMs subjected to varying matrix rigidity (Young’s modulus of 1, 10, and 100 kPa) on fibronectin-coated polyacrylamide (PA) gel as a culture platform in the absence or presence of IL4/13. The 2-dimensional (2D) cellular areas increased significantly with increasing matrix rigidity (1 kPa < 10 kPa < 100 kPa) (**Fig. 1A and Fig. S1A**). Notably, these morphological changes were independent of IL4/13 treatment (**Fig. 1A**), indicating that matrix rigidity plays a pivotal role in shaping cellular morphology – a well-recognized mechanotype that reflects the mechano-activated/-primed cell state ^11^. Similar morphological variations dependent on matrix rigidity were observed in RAW264.7 (a mouse macrophage cell line) and THP-1 (a human monocyte/macrophage cell line) cells (**Fig. S1B, C**).

**Figure 1.**
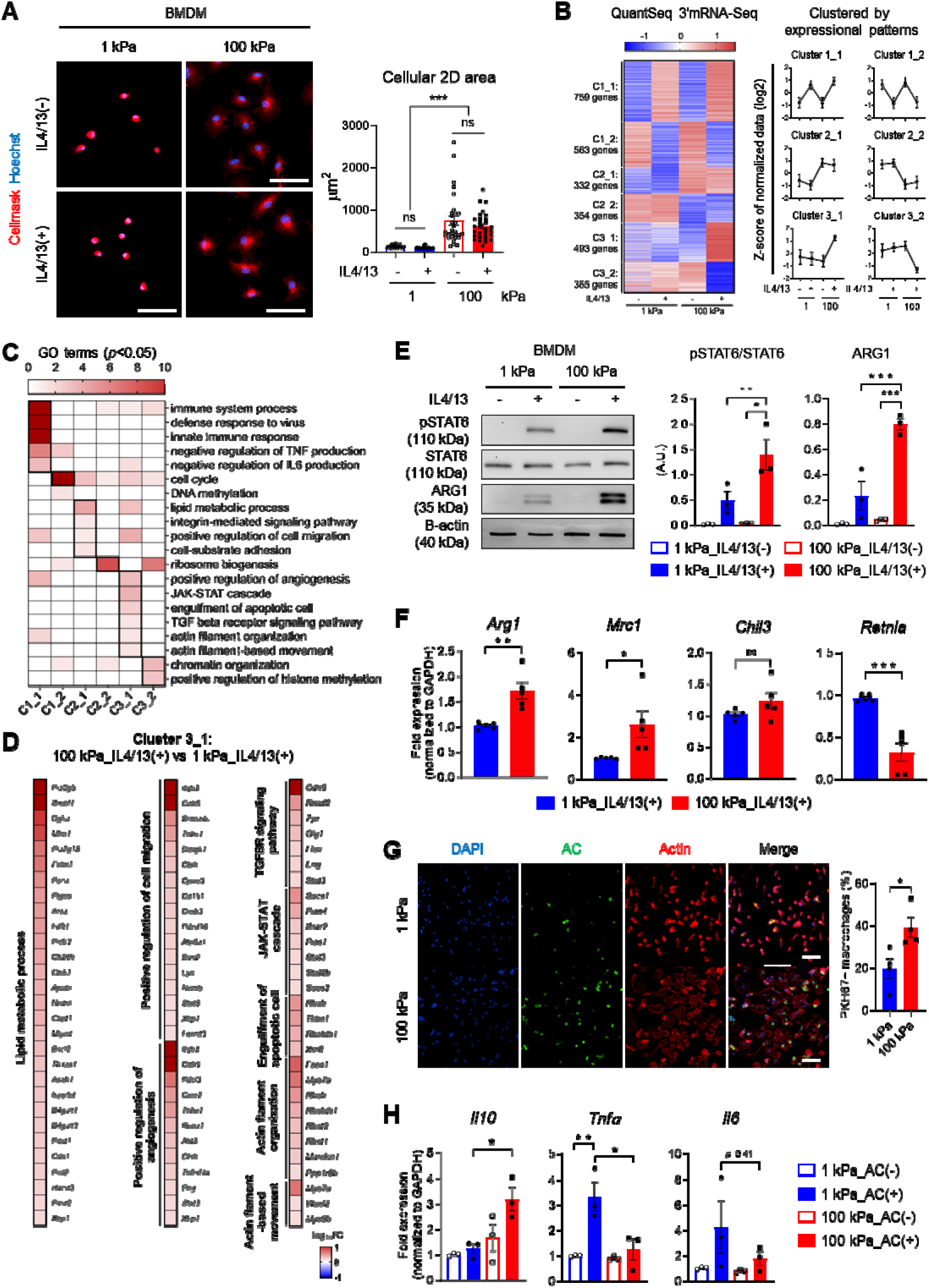
High matrix rigidity promotes IL4/13-mediated M2 macrophage activation. (**A**) Representative images of CellMask (red) and Hoechst-33342 (blue) staining in mouse bone marrow-derived macrophages (BMDMs) cultured on either a 1 kPa or 100 kPa polyacrylamide (PA) gel with or without 20 ng/mL IL4/13 treatment (scale bars = 50 µm). The 2-dimensional (2D) cellular surface area was quantified (*n* = 30 cells/group) and presented as mean ± SEM (*** *p* < 0.001; Kruskal-Wallis test). ns, not significant. **(B)** Heatmap depicting differentially expressed genes (DEGs) from QuantSeq 3’ mRNA-Seq analysis (|fold change (FC)|≥1.5). DEGs are categorized into six clusters based on expression patterns: up/downregulated by IL4/13 treatment regardless of substrate rigidity (C1_1/C1_2), up/downregulated by matrix rigidity regardless of IL4/13 treatment (C2_1/C2_2), and up/downregulated only in BMDMs cultured on 100 kPa with IL4/13 treatment compared to the other groups (C3_1/C3_2). Line plots show the mean ± standard deviation of Z-score of normalized log_2_ values from QuantSeq 3’ mRNA-Seq dataset. **(C)** Heatmap representing 20 biological processes in each cluster, with –log_10_ raw binomial *p*-values determined by DAVID Gene Ontology (GO) term enrichment analysis. **(D)** Heat map showing log_10_-converted FC ratios of Cluster 3_1 DEGs associated with lipid metabolic process, positive regulation of cell migration, positive regulation of angiogenesis, TGFBR signaling pathway, JAK-STAT cascade, engulfment of apoptotic cell, actin filament organization, and actin filament-based movement in the 100kPa_IL4/13(+) condition compared to 1kPa_IL4/13(+) condition. **(E)** Representative Western blots and quantification of phospho-STAT6 (pSTAT6), STAT6, ARG1, and B-actin in BMDMs cultured on 1 kPa or 100 kPa PA gels with or without IL14/13 (20 ng/mL) treatment for 12 hours. Results are presented as mean ± SEM (*n* = 3 replicates; * *p* < 0.05, ** *p* < 0.005, and *** *p* < 0.001; one-way ANOVA). A. U.: Arbitrary Unit. **(F)** qRT-PCR analysis of M2-associated genes (*Arg1*, *Mrc1*, *Chil3*, and *Retnla*) in BMDMs cultured on 1 kPa or 100 kPa PA gels with IL4/13 treatment (*n* = 5 replicates; * *p* < 0.05, ** *p* < 0.005, and *** *p* < 0.001; unpaired Student *t*-test). ns, not significant. **(G)** Representative images of phalloidin (red)– and DAPI (blue)-stained BMDMs on 1 kPa or 100 kPa PA gels. Efferocytotic BMDMs were identified by co-localization with PKH67-labelled apoptotic Jurkat cells (AC, green), and percentage of PKH67+ BMDMs in total BMDMs was quantified and presented as mean ± SEM (*n* = 4 replicates; \**p* < 0.05; unpaired Student *t*-test). **(H)** qRT-PCR analysis of *Il10*, *Tnf*, and *Il6* in BMDMs cultured on 1 kPa or 100 kPa gels with or without ACs. (*n* = 3 replicates; * *p* < 0.05, and ** *p* < 0.005; one-way ANOVA).

Next we conducted global mRNA expression profiling using QuantSeq 3’ mRNA-Seq analysis on BMDMs cultured with or without IL4/13 treatment on compliant (1 kPa) or rigid (100 kPa) substrates. A total of 2,866 mRNAs were identified to be differentially expressed (|FC|≥1.5, normalized data (log2)≥4) and classified into six clusters based on expression patterns across the different conditions (**Fig. 1B**): up/downregulated by IL4/13 treatment regardless of matrix rigidity (Cluster 1_1/Cluster 1_2), up/downregulated by matrix rigidity regardless of IL4/13 treatment (Cluster 2_1/Cluster 2_2), and up/downregulated only in BMDMs cultured on 100 kPa with IL4/13 treatment compared to the other groups (Cluster 3_1/Cluster 3_2). Gene Ontology (GO) enrichment analysis revealed that mRNAs related to immune system process, defense response to virus, innate immune response, and negative regulation of TNF and IL6 production were upregulated mainly by IL4/13 treatment (**Fig. 1C**, Cluster 1_1). Lipid metabolic process, integrin-mediated signaling pathway, positive regulation of cell migration, and cell-substrate adhesion, which are mechanotransducive pathways, were engaged by matrix rigidity in cluster 2_1. Notably, cluster 3_1 genes were enriched in pathways linked to M2 macrophage features, including positive regulation of angiogenesis, JAK-STAT cascade, engulfment of apoptotic cell, and TGF-β receptor signaling pathway (**Fig. 1C, D**), indicating a potential cooperative role of mechano-chemical signals in promoting M2 activation. In addition, actin filament organization and actin filament-based movement, chromatin organization, and positive regulation of histone methylation were the pathways that are influenced by interplays between IL4/13 (biochemical cue) and substrate rigidity (mechanical cue) (**Fig. 1C, D**).

Given the crucial role of the IL4/13-JAK-STAT6 pathway in M2 macrophage activation ^17^, the phosphorylation state of STAT6 was assessed by immunoblotting. Notably, STAT6 phosphorylation was significantly enhanced on rigid substrates with minimal-to-optimal concentrations of IL4/13 (**Fig. 1E and Fig. S2A**), resulting in elevated expression levels of STAT6-transcribed M2 markers (*Arg1*, *Mrc1*) on 100 kPa compared to 1 kPa (**Fig. 1F**). We further observed that the expression of arginase 1 (ARG1) expression at the protein level was augmented with increasing matrix rigidity from 1 kPa to 10, 23, and 100 kPa (**Fig. 1E and Fig. S2B**). Interestingly, while expression levels of pSTAT6 during an early period of IL4/13 treatment (15 min, 1 h, 3 h) were comparable between 1 kPa and 100 kPa, the pSTAT6 level was markedly higher on 100 kPa compared to 1 kPa at the later time point (12 h) when ARG1 expression was evident (**Fig. S2C**). This indicates sustained activation of the IL4/13-JAK-STAT6 pathway under rigid substrate conditions. This mechano-chemical activation of M2 macrophages also occurred in RAW264.7 and THP-1 cells, which was evidenced by the upregulations of pSTAT6 and ARG1 levels on rigid substrates in the presence of IL4/13 (**Fig. S3**), and in BMDMs through IL10-STAT3 pathway ^18,19^ (**Fig. S4**), implying that matrix mechanics interplay with biochemical signals pervasively in the activation of M2.

To assess the functional consequence of elevated M2 markers on rigid substrates, we examined the M2 macrophage ability of phagocytic clearance of apoptotic cells (ACs), called efferocytosis ^20,21^, under different rigidity conditions. We found that BMDMs on rigid matrix exhibited enhanced efferocytotic activity compared to those on compliant one (**Fig. 1G**). qRT-PCR analysis indicated increased expression of the anti-inflammatory cytokine *Il10* and reduced expression of pro-inflammatory cytokines *Tnfα* and *Il6* on rigid matrix (**Fig. 1H**), suggesting that higher matrix rigidity may bolster efferocytosis, contributing to inflammation resolution and tissue regeneration. Taken together, these findings indicate that the rigidity of cell-adhesive substrates work cooperatively with immunomodulatory cytokines like IL4/13 and IL10 to reinforce the activation of M2 macrophages, holding implications for modulating inflammation and tissue repair.

### Matrix rigidity induces chromatin reorganization, marked by downregulation of H3K9me2/3 with liberated LADs at the nuclear periphery

Unlike *Arg1* and *Mrc1*, other M2 markers, such as *Chil3* and *Retnla*, were unaffected or downregulated by increased matrix rigidity, although they were upregulated by IL4/13 treatment in both rigidity conditions (**Fig. 1F and Fig. S5**). These distinct responses of gene regulation to matrix rigidity prompted us to investigate potential selective effects on genomic regions through chromatin reorganization and physical interactions among promoters, enhancers, insulators, and chromatin-binding factors within accessible chromatin areas ^22–25^. Studies have revealed that certain mechanical cues can regulate epigenetic pathways that are known to be important in governing macrophage behavior ^26^. We investigated whether substrate rigidity alters di-or tri-methylated histone H3 at lysine 9 (H3K9me2/3) levels, indicative of heterochromatin and transcriptional silencing, since H3K9me2/3 is shown to play an important role in macrophage polarization ^27–29^. Analysis of H3K9me2/3 expression profiles across the nuclear XY axis revealed distinct localization patterns. BMDMs on compliant (1 kPa) substrates exhibited concentrated H3K9me2/3 at the nuclear rim (**Fig. 2A**). In contrast, upon rigid (100 kPa) substrates the peripheral expression of H3K9me2/3 was markedly reduced (**Fig. 2B**), and the overall H3K9me2/3 levels were also significantly downregulated (**Fig. 2C**). Importantly, IL4/13 treatment did not alter the H3K9me2/3 levels in either rigidity condition (**Fig. 2A-C**), suggesting matrix rigidity mechano-primes the epigenetic landscape of macrophages, especially H3K9me2/3 levels.

**Figure 2.**
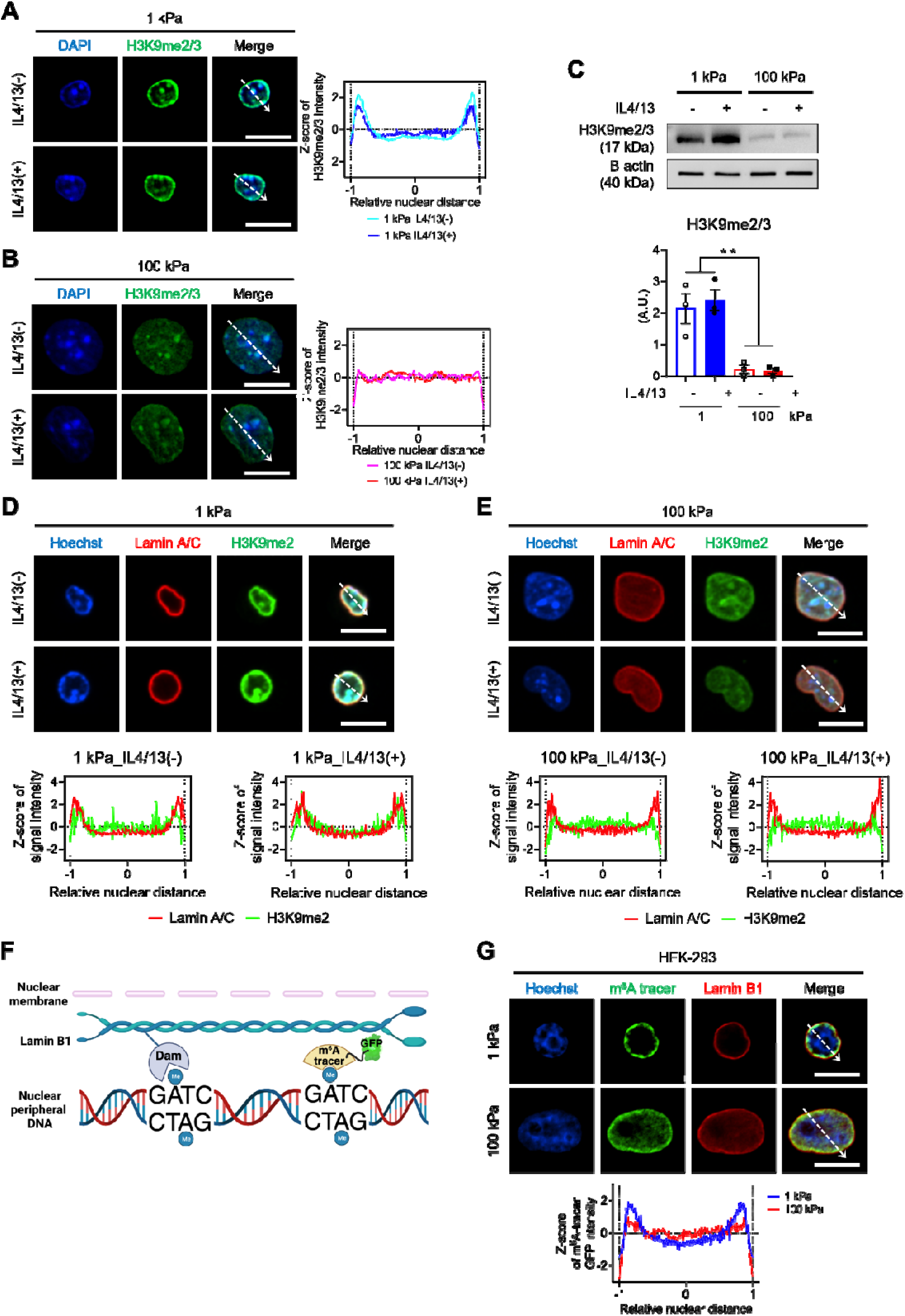
Lamina-associated domains (LADs) are reduced in macrophages by sensing high matrix rigidity. (**A, B**) Representative immunostaining images of DAPI (blue) and H3K9me2/3 (green) in BMDMs cultured on either a 1 kPa **(A)** or 100 kPa **(B)** PA gel with or without IL4/13 treatment (scale bars = 10 µm). The mean Z-scores of normalized H3K9me2/3 signal intensity across the relative nuclear distance indicated by dashed arrows (*n* = 30 for 1 kPa_IL4/13(-), *n* = 30 for 1 kPa_IL4/13(+), *n* = 31 for 100 kPa_IL4/13(-), *n* = 29 for 100 kPa_IL4/13(+)) are shown. 0 represents the center of the nucleus, and ± 1 represents the nuclear periphery. **(C)** Representative Western blots and quantification of H3K9me2/3 and B-actin in BMDMs cultured on 1 kPa or 100 kPa PA gels with or without IL4/13 treatment. Results are presented as mean ± SEM (*n* = 3 replicates; ** *p* < 0.005; one-way ANOVA). **(D, E)** Representative co-immunostaining images of Lamin A/C (red), H3K9me2 (green), and Hoechst-33342 (blue) in BMDMs cultured on either a 1 kPa **(D)** or 100 kPa **(E)** PA gel with or without IL4/13 treatment (scale bars = 10 µm). Dashed arrows indicate positions of the line signal intensity profiles. **(F)** A schematic image illustrating the DNA adenine methyltransferase identification (DamID) technique. In the presence of DNA near the nuclear lamina, Lamin B1-linked Dam induces adenine methylation, enabling the subsequent identification of N^6^-methyladenosine through m^6^A tracer conjugated with GFP. **(G)** Representative images of Hoechst-33342 (blue), m^6^A tracer (GFP, green), and Lamin B1 immunostaining (Red) in HEK-293 cells cultured on 1 kPa or 100 kPa PA gels and co-transfected with the DD-DamWT-LMNB1-IRES2-mCherry plasmid and m^6^A-Tracer-NES plasmid for 48 h. The mean Z-scores of normalized m^6^A tracer GFP intensity are presented across the relative nuclear distance, as indicated by the dashed arrows (*n* = 30 nuclei per each condition).

The large heterochromatin compartment located in the nuclear lamina is called lamina-associated domain (LAD) ^22,23^. Most genes in LADs are transcriptionally repressed ^24,25^. Through co-immunostaining for H3K9me2 and Lamin A/C, we validated the reduction of heterochromatin mark within the nuclear lamina upon increased rigidity, where H3K9me2-marked heterochromatin appeared highly associated with nuclear lamina in the form of LADs (**Fig. 2D, E**). This observation suggests that certain genomic loci were loosened from the nuclear periphery of BMDMs on rigid substrates. Consistent patterns of LAD abundance between rigid and compliant substrates were also observed in RAW264.7 and THP-1 cells (**Fig. S6**). Furthermore, IL4/13 treatment had no impact on LAD abundance (**Fig. 2D, E**). To further substantiate that substrate rigidity induces changes in LADs, we utilized the DNA adenine methyltransferase identification (DamID) technique to fluorescently label Dam-mediated N^6^-methylated adenosine (m^6^A) residues within DNA regions that interact the nuclear lamina ^30,31^ (**Fig. 2F**). HEK-293 cells on rigid substrates demonstrated fewer LADs compared to cells on compliant substrates (**Fig. 2G**). These findings indicate that LADs undergo reorganization in response to matrix rigidity, possibly influencing chromatin accessibility and thereby modulating the transcriptional expression of key genes involved in M2 macrophage activation.

### ATAC-seq reveals enhanced chromatin accessibility of M2-associated gene promoters, reorganized upon mechano-chemical cues

We then performed assays for transposase-accessible chromatin with high-throughput sequencing (ATAC-seq) for BMDMs cultured either on compliant or rigid substrates in the absence or presence of IL4/13, to assess if chromatin accessibility in upstream/promoter of M2-associated genes is distinctly reorganized by mechano-chemical stimuli. The calculation of Transcription starting site (TSS) enrichment scores revealed that chromatin-accessible regions are enriched at TSSs and aligned fragment length distribution of all samples indicated that the ATAC-seq data is of high quality (**Fig. S7A, B**). Chromatin-accessible regions were distributed throughout the genome similarly between different conditions, and approximately 30% of them were upstream/promoters (∼3 kb from TSS) (**Fig. S7C**). We discovered 2,203 differentially decondensed or condensed regions in upstream/promoter/overlap to 5’ sequences (|FC|≥1.5, normalized data (log2)≥1; distinct regulatory elements, DREs) accounting for 21.4% of all DREs **(Fig. S7D)**, and classified them into six distinct clusters based on chromatin accessibility patterns across 4 combination conditions (**Fig. 3A**): decondensed/condensed by IL4/13 treatment regardless of matrix rigidity (Cluster 1_1/Cluster 1_2), decondensed/condensed by matrix rigidity regardless of IL4/13 treatment (Cluster 2_1/Cluster 2_2), and decondensed/condensed only in BMDMs cultured on 100 kPa with IL4/13 treatment compared to the other groups (Cluster 3_1/Cluster 3_2). This clustering showed an apparent chromatin reorganization of genes, which was promoted by either matrix rigidity, IL4/13, or both. Functional annotations of genes with de-condensed chromatin regions (clusters C1_1, C2_1, C3_1) showed that M2-featured functions associated with fatty acid metabolic process ^32^, positive regulation of cell migration and angiogenesis ^33^, and positive regulation of STAT cascade ^34^ were enriched during alternative activation of macrophages (**Fig. 3B**).

**Figure 3.**
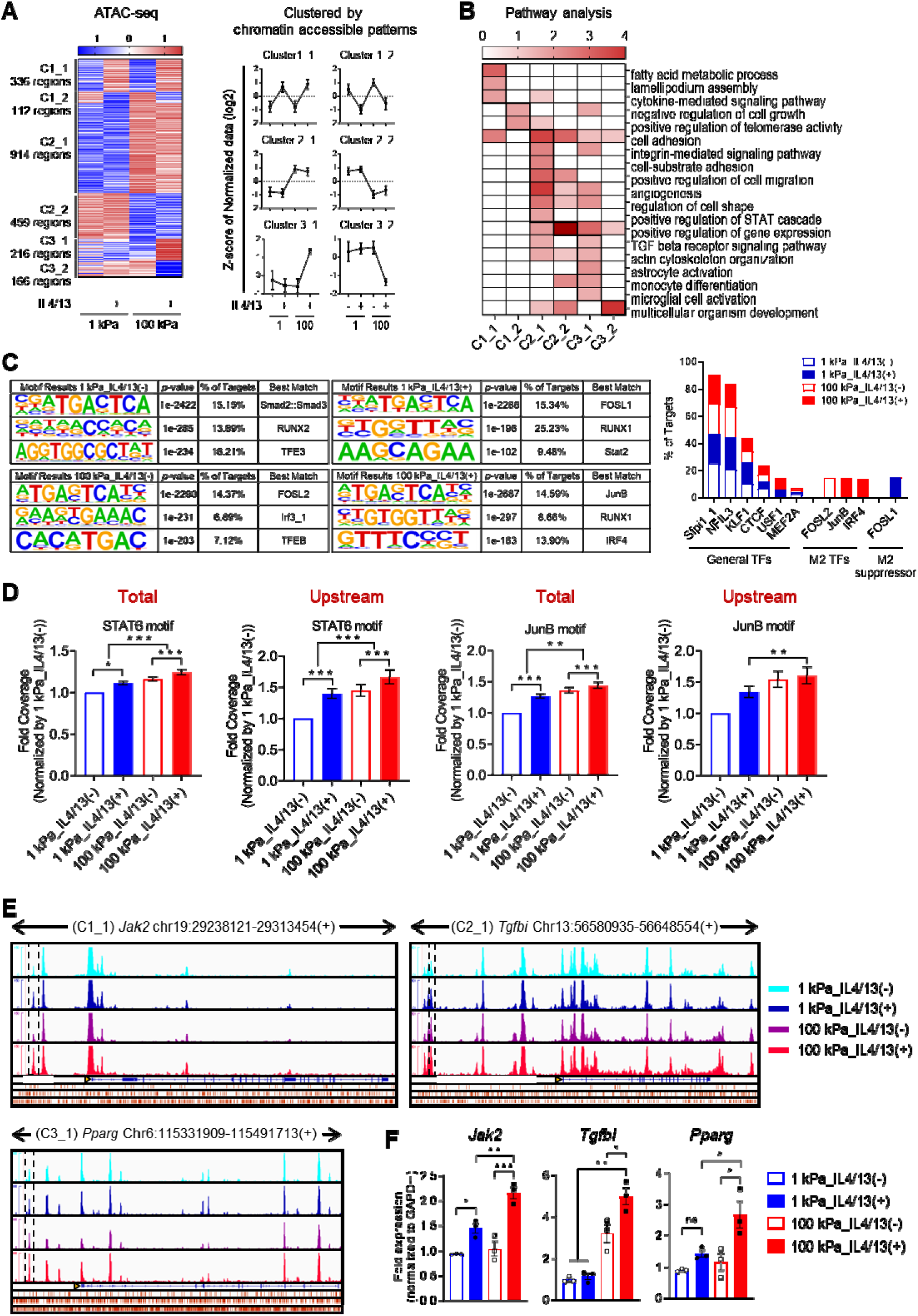
Chromatin accessibility changes in response to mechano-chemical stimuli. (**A**) Heatmap illustrating distinct regulatory elements (DREs) within promoter and upstream regions from ATAC-seq analysis (|FC|≥1.5). DEGs are categorized into six clusters based on chromatin accessible patterns: open/closed by IL4/13 treatment regardless of matrix rigidity (C1_1/C1_2), open/closed by matrix rigidity regardless of IL4/13 treatment (C2_1/C2_2), and open/closed only in BMDMs cultured on 100 kPa with IL4/13 treatment compared to the other groups (C3_1/C3_2). Line plots depict the mean ± standard deviation of Z-score of normalized log_2_ values from ATAC-seq dataset. **(B)** Heatmap displaying 19 biological processes in each cluster presented as –log_10_ raw binomial *p*-values, calculated by DAVID GO term enrichment analysis. **(C)** Representative *de novo* transcription factor (TF) binding motifs discovered using Homer software in four different BMDM culture conditions (1 kPa_IL4/13(-), 1 kPa_IL4/13(+), 100 kPa_IL4/13(-), and 100 kPa_IL4/13(+)). Bar graph indicates the percentage of targets of binding motifs of general TFs (Sfpi1_1, NFIL3, KLF1, CTCF, USF1, MEF2A), M2 macrophage activation-associated TFs (FOSL2, JunB, IRF4), and a TF suppressing M2-associated gene expression (FOSL1). **(D)** The bar plots depicting the fold change of log_2_-normalized coverage levels on TF binding motifs in both the entire genome regions and the upstream regions (* *p* < 0.05, ** *p* < 0.005, *** *p* < 0.001; Kruskal-Wallis test). **(E)** Normalized ATAC-seq tracks of samples across four distinct BMDM culture conditions along the gene locus in C1_1, C2_1, and C3_1 using Integrative Genomics Viewer (IGV). DREs are delineated with dashed boxes, and these elements are situated in the promoter/upstream regions relative to the transcription start site (indicated by a yellow arrowhead). Binding sites corresponding to M2 macrophage-associated TFs, including STAT6 (top), IRF4 (middle), and JunB (bottom), are denoted with red bars. **(F)** qRT-PCR analysis of *Jak2*, *Tgfbi*, and *Pparg* in BMDMs cultured on 1 kPa or 100 kPa PA gels with or without IL4/13 treatment (*n* = 3 replicates; * *p* < 0.05, ** *p* < 0.005, and *** *p* < 0.001; one-way ANOVA).

Notably, Homer *de novo* TF binding motif discovery predicted that a couple of TFs that promote alternative activation of macrophages, JunB and IRF4 ^35,36^, can bind to chromatin-accessible regions exclusively in the condition of 100 kPa with IL4/13 (*p*<0.05) (**Fig. 3C**), explaining partly how M2 polarization was promoted when macrophages were exposed to both mechanical and chemical signals compared to when only either one of them was present. On the contrary, FOSL1 that belongs to AP-1 family downregulating *Arg1* transcription by directly binding to *Arg1* promoter regions ^37^ was shown to have exclusive, highest frequency of binding motifs in promoter regions in the 1 kPa plus IL4/13 condition (**Fig. 3C**). General TFs, including Sfpii1_1 (PU.1), KLF1, CTCF, NFIL3, and MEF2A, were predicted to have their binding motifs with comparable frequencies among 4 conditions. The top 10 TFs predicted to have binding motifs in chromatin-accessible regions of each condition were shown in **Fig. S8**. Furthermore, an analysis of chromatin openness in regions containing TF binding motifs revealed that IL4/13 treatment and increased matrix rigidity led to increased chromatin decondensation in STAT6 binding motifs in the whole genomic regions (**Fig. 3D**). Especially, in the upstream/promoter regions, the binding motifs of STAT6 and JunB (MA0520.1 and MA0490.2, respectively) exhibited significant decondensation due to cooperative effect of both IL4/13 treatment and increased matrix rigidity (**Fig. 3D**).

We validated this by confirming that several chromatin-accessible regions, the upstream/promoter regions of genes associated with the M2 phenotype, displayed higher peaks under IL4/13-treated conditions (**Fig. 3E** and **Fig. S9A**) and in the 100 kPa rigidity conditions (**Fig. 3E** and **Fig. S10A**). Also, the chromatin regions that became condensed due to IL4/13 treatment or increased matrix rigidity are presented in **Fig. S9B** and **Fig. S10B**, respectively. Specifically, the upstream/promoter regions of classical M2 markers, including *Jak2* (Chr19:29239500-29239900) and *Arg1* (Chr10:24980670-24981070), were part of the C1_1 cluster of which chromatin accessibility increased in response to IL4/13 treatment (**Fig. 3E** and **Fig. S9A**). The C2_1 cluster contained chromatin regions associated with *Tgfbi* (Chr13:56582628-56583028), a gene strongly linked to tissue remodeling and repair ^38,39^ (**Fig. 3E**). Interestingly, the upstream/promoter regions of M2-associated genes, such as *Pparg* (Chr6:115334439-115334839), *Tgm2* (Chr2:158159895-15816029, 158161660-

158162060), and *Irf4* (Chr13:30738430-30738830), which possess STAT6, IRF4, or JunB binding motifs, showed the highest peaks under the 100 kPa plus IL4/13 treatment condition (**Fig. 3E** and **Fig. S11A**). In contrast, M1-associated genes like *Il6* (Chr5:29972808-29973208) and *Tlr2* (Chr3:83844786-83845186) ^40^, which have the FOSL1 binding motifs, displayed the lowest peaks under the same condition (**Fig. 3E** and **Fig. S11B**). Importantly, alterations in chromatin accessibility induced by cytokines and/or mechanical stress correlated well with changes in the transcriptional levels of the corresponding genes (**Fig. 3F** and **Fig. S11C**). These findings suggest that on rigid substrates, chromatin organization may be structured to promote M2 macrophage activation while inhibiting M1 macrophage activation.

### Formation of mechanosensitive podosome-like actin structure and actomyosin contractility are responsible for M2 macrophage activation

Cellular sensing of mechanical signals from their microenvironment leads to cytoskeletal rearrangements through integrin-mediated signaling pathways, resulting in the formation of actin-rich structures, such as podosomes in macrophages ^41^. Podosomes are columnar actin structures that serve functions in adhesion, migration, proteolysis, and mechanosensing ^42^. To investigate the role of mechanosensing in M2 macrophage activation, we examined the impact of matrix rigidity on actin cytoskeletal arrangements in BMDMs. On relatively compliant substrates (1 kPa), BMDMs primarily exhibited cortical actin arrangements, whereas a substantial increase in podosome-like actin dot structures was observed on rigid substrates (10, 100 kPa) (**Fig. 4A**). The densities of actin dots were consistent across substrates, likely due to variations in cytoplasmic area (**Fig. 4A**). These actin dot structures were encircled by a ring composed of integrin-associated proteins, such as vinculin (**Fig. 4B**), characteristic of podosomes ^42–45^. IL4/13 treatment did not influence the cytoskeletal rearrangements.

**Figure 4.**
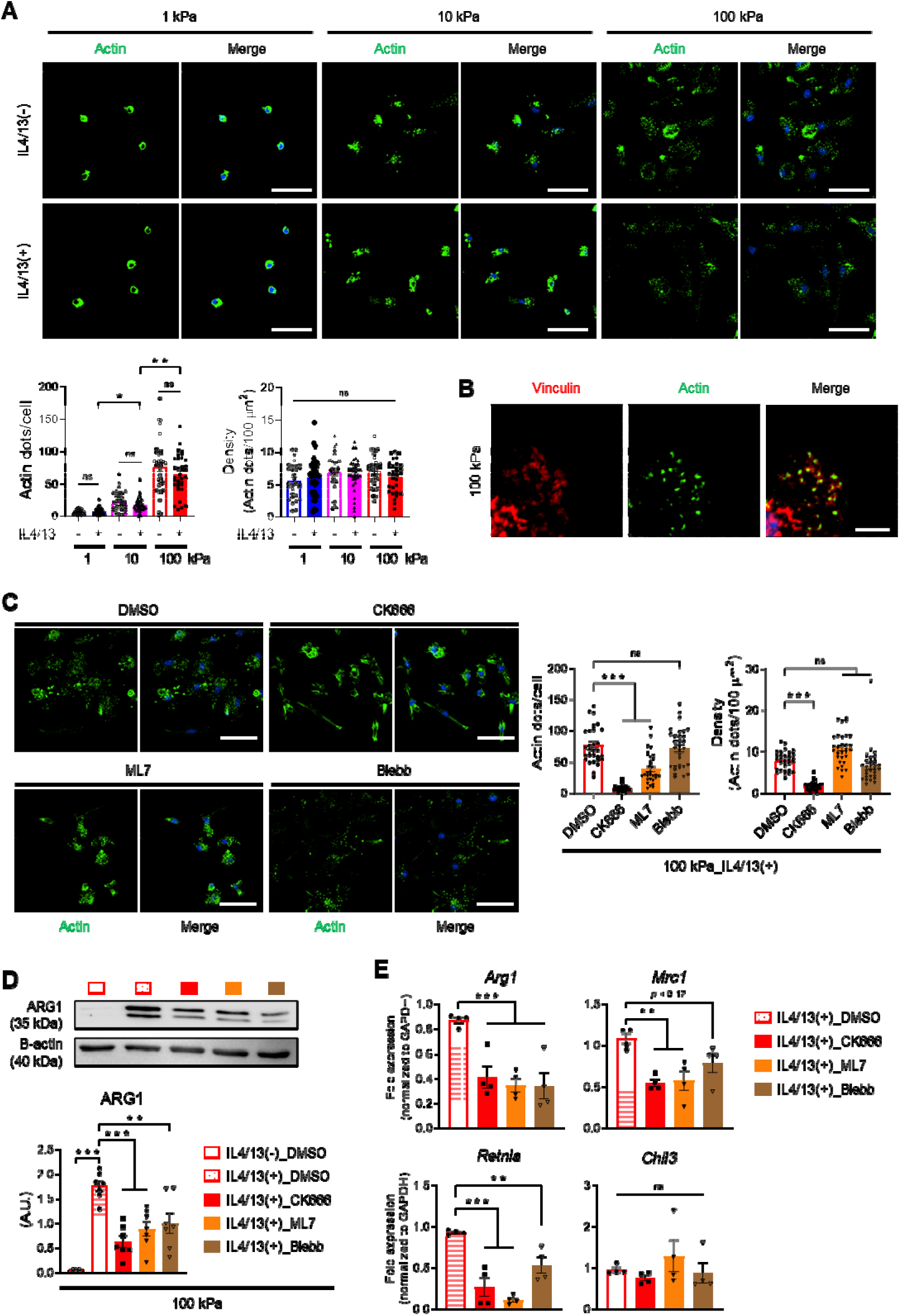
Podosome-like actin dot structure formation and actomyosin contractility responsible for the acceleration of M2 macrophage activation. (**A**) Representative images of phalloidin (green) and DAPI (blue)-stained BMDMs cultured on 1 kPa, 10 kPa, or 100 kPa PA gels with or without IL4/13 treatment (scale bars = 50 µm). The number of actin dots per cell and density (number of actin dots/100 µm^2^) are presented as mean ± SEM (*n* = 30 cells/group; * *p* < 0.05 and *** *p* < 0.001; one-way ANOVA). ns, not significant. **(B)** Representative immunostaining images of Vinculin (red), phalloidin (green), and DAPI (blue) in BMDMs cultured on 100 kPa PA gel 8 h after seeding in the absence of IL4/13 (scale bar = 5 µm). **(C)** Representative images of phalloidin (green) and DAPI (blue)-stained BMDMs cultured on 100 kPa PA gel with DMSO, CK666 (200 µM), ML7 (25 µM), or Blebbistatin (Blebb, 50 µM) in the presence of IL4/13 (scale bars = 50 µm). The number of actin dots per cell and density (number of actin dots/100 µm^2^) are shown as mean ± SEM (*n* = 30 cells/group; *** *p* <0.001; Kruskal Wallis test). ns, not significant. **(D)** Representative Western blots and quantification of ARG1 and B-actin in BMDMs under different culture conditions (*n* = 5 for IL4/13(-)_DMSO, and *n* = 7 for IL4/13(+)_DMSO, IL4/13(+)_CK666, IL4/13(+)_ML7, and IL4/13(+)_Blebb). Results are presented as mean ± SEM (** *p* < 0.005 and *** *p* < 0.001; one-way ANOVA). **(E)** qRT-PCR analysis of M2 macrophage-associated genes (*Arg1*, *Mrc1*, *Chil3*, and *Retnla*) in BMDMs cultured on 100 kPa treated with DMSO, CK666 (200 µM), ML7 (25 µM), or Blebb (50 µM) in the presence of IL4/13 (*n* = 4 replicates; * *p* < 0.05, ** *p* < 0.005, and *** *p* < 0.001; one-way ANOVA or Kruskal-Wallis test). ns, not significant.

To ascertain the role of podosome-like actin structures in macrophage activation, BMDMs on 100 kPa rigidity were treated with inhibitors CK666 (inhibitor of Arp2/3-mediated actin polymerization), ML7 (myosin light chain kinase inhibitor), and blebbistatin (selective inhibitor of ATPase activity of myosin II) to destruct actin structures or disrupt actin-mediated traction forces. CK666 treatment resulted in the loss of podosome-like actin dot structures and concurrent downregulation of classical M2 macrophage markers, including *Arg1*, *Mrc1*, and *Retnla*, at mRNA or protein levels (**Fig. 4C-E**). While actin dot structures were largely preserved following treatment with inhibitors targeting actomyosin contractility (ML7 and blebbistatin) to the level comparable to DMSO control, expressions of M2-associated genes were downregulated (**Fig. 4C-E**). These findings underscore the significance of podosome-like actin structures and actomyosin-mediated force transmission in expediting M2 macrophage activation.

### Matrix rigidity induces nuclear deformation and surface tension increase through cytoskeletal-mediated nuclear mechanosensing

Podosomes are recognized for generating increased traction forces with escalating substrate rigidity^46^. Our previous work ^11^ demonstrated that nuclear import of STAT6 is enhanced in BMDMs under high matrix rigidity conditions through F-actin-mediated nuclear mechanosensing that is represented by the increased nuclear flattening. Here we sought to validate altered nuclear morphologies across different rigidity and cytokine conditions and confirm the role of cytoskeletal-mediated force transmission in promoting nuclear import of pSTAT6. Measurement of nuclear 2D projection area, flattening, and 3D surface area in BMDMs (**Fig. S12**) revealed significant increases on rigid substrates compared to compliant ones, irrespective of IL4/13 treatment (**Fig. 5A, B and S13A**). These nuclear parameters were substantially reduced following treatment with cytoskeletal inhibitors CK666, ML7, and blebbistatin (**Fig. 5C, D and Fig. S13B**), demonstrating the nuclear deformation is governed by the cytoskeletal mechanics.

**Figure 5.**
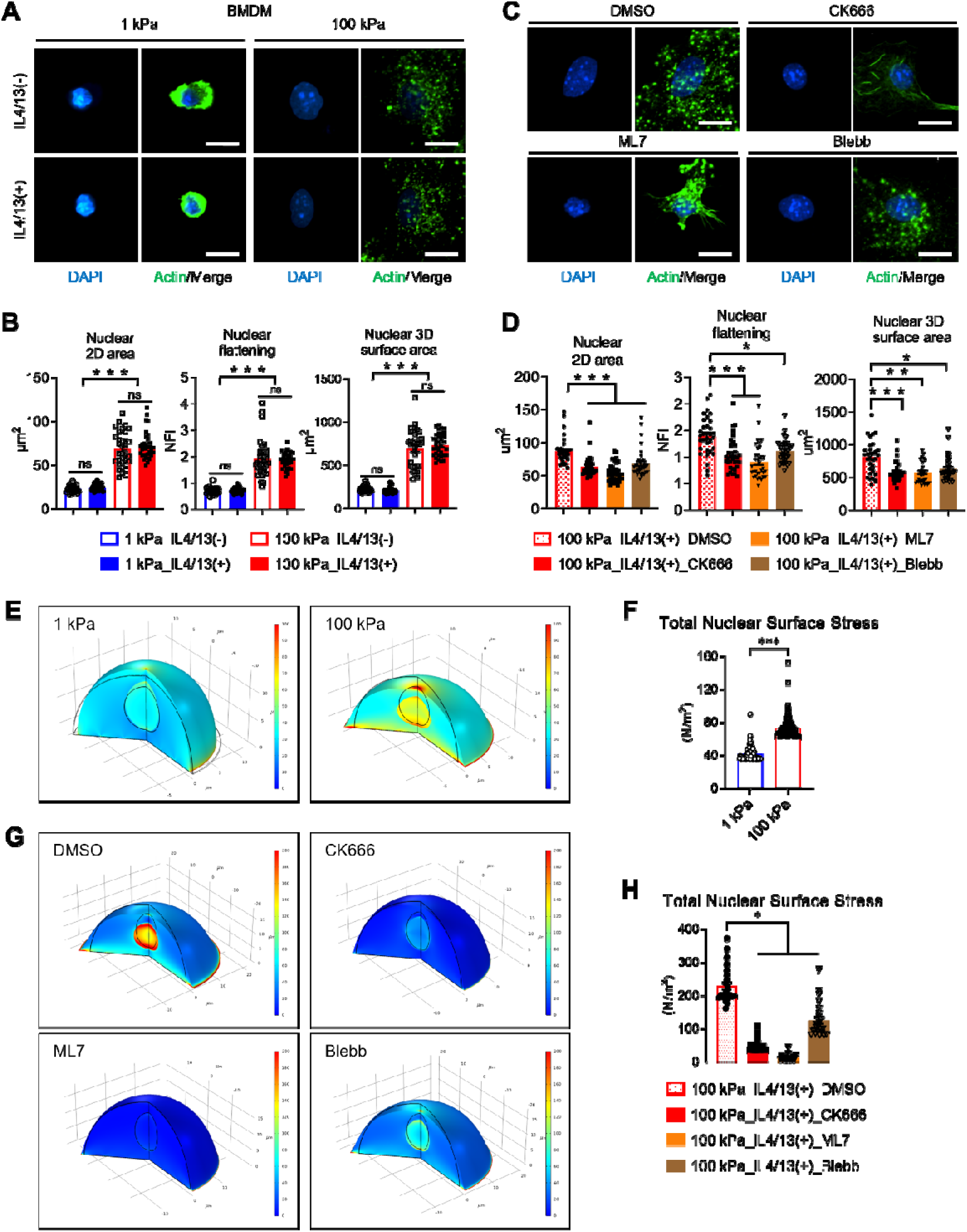
Nuclear deformation and increased surface tension on rigid matrices via cytoskeletal-mediated nuclear mechanosensing. (**A**) Representative images of phalloidin (green) and DAPI (blue)-stained BMDMs cultured on 1 kPa or 100 kPa PA gels with or without IL4/13 treatment (scale bars = 10 µm). **(B)** Measurements of nuclear 2D area (µm^2^), nuclear flattening index (NFI), and nuclear 3D surface area (µm^2^) of BMDMs on 1 kPa or 100 kPa PA gels with or without IL4/13 treatment. Results are presented as mean ± SEM (*n* = more than 30 cells/group; *** *p* < 0.001; one-way ANOVA or Kruskal-Wallis test). ns, not significant. **(C, D)** Representative images **(C)** and measurements of nuclear 2D area, NFI, and nuclear 3D surface area **(D)** of BMDMs on 100 kPa PA gel with DMSO, CK666 (200 µM), ML7 (25 µM), or Blebb (50 µM) treatment in the presence of IL4/13 (scale bars = 10 µm). Results are presented as mean ± SEM (*n* = more than 30 cells/group; * *p* < 0.05 and \*\*\**p* < 0.001; Kruskal-Wallis test). ns, not significant. **(E)** 3D reconstructed views illustrating the deformation and stress distribution within the cell anchored to a compliant (left) or rigid (right) substrate. Cell deformation was simulated through horizontal stretching, and boundary loads were applied to the cytoplasm. The black solid curves indicate the nucleus and cell boundary before deformation, with the nucleus positioned at the center of the hemispherical cell model. Stress distribution after deformation was color-coded. **(F)** The mean ± SEM of total von-Mises stress applied to the nuclear surface of cells on rigid substrates was significantly higher than that of cells on compliant substrates (*** *p* < 0.001; Mann-Whitney test). **(G)** Deformation and stress distribution within the cell treated with DMSO, CK666, ML7, or Blebb in the rigid substrate condition. **(H)** The mean ± SEM of total nuclear surface stress of the cells treated with cytoskeletal inhibitors were significantly reduced compared to DMSO-treated cells (*\* p* < 0.05; Kruskal-Wallis test).

We further employed a computational model to simulate the nuclear deformation and mechanical stress of BMDMs dependent on matrix rigidity. In our simulation, we initially represented a model cell adhering to either a compliant or rigid substrate as a hemisphere with a radius of 12 µm. Within the cell, the nucleus was depicted as an ellipsoidal shape with a vertical radius of 4 µm and a horizontal radius of 3.5 µm. To describe the mechanical properties of the cell, we utilized a linear elastic material with a viscoelastic component, a generalized Maxwell model. The cell’s mechanical characteristics are outlined in **Table S1**. In our model, we simulated nuclear deformation by applying pulling forces to the bottom of the cell adhering to the deformable substrates. This resulted in the generation of pressure by the cell membrane, which was assumed to connect to the nucleus via the intracellular cytoskeletons. The results demonstrated nuclear deformation and stress distribution within the cell, depending on whether it adhered to a compliant or rigid substrate (**Fig. 5E and Fig. S14A**). On the compliant substrate, the nucleus transformed into a standing egg-like shape, while on the rigid substrate, it appeared as a lying egg at the bottom of the cell. Notably, the magnitude of von-Mises stresses on the nucleus was observed to be higher on its surface compared to its center. Also, quantification of von-Mises stress on the nuclear surface revealed a gradual decrease from the center to the boundary of the cell (**Fig. S14B**). The total von-Mises stress applied to the nuclear surface of a cell placed on a rigid substrate was significantly higher (about 1.7-fold) than that of a cell on a compliant substrate (**Fig. 5F**). On the other hand, the total nuclear surface stresses were extensively reduced when actomyosin was disrupted by treatment with CK666, ML7, and blebbistatin (**Fig. 5G, H, and Fig. S14C**), in line with experimental observations in nuclear morphological changes (**Fig. 5D and Fig. S13B**). These results implicate that matrix rigidity positively modulates the physical deformation of nucleus via formation of podosome-like actin cytoskeletal structure and actomyosin contractility.

### Deformed nucleus induces pore enlargement, facilitating pSTAT6 nuclear translocation

Along with the nuclear deformation, immunostaining and nuclear/cytoplasmic protein fractionation consistently demonstrated enhanced nuclear import of pSTAT6 in IL4/13-treated macrophages on rigid substates (**Fig. 6A-C**). The nuclear-to-cytoplasmic ratio of pSTAT6 levels correlated positively with nuclear flattening and 3D surface area in IL4/13-treated THP-1 cells (**Fig. 6B**). Moreover, pSTAT6 expression within the nucleus was significantly diminished upon actomyosin inhibitor treatment in IL4/13-treated BMDMs (**Fig. 6D**), along with the decrease of nuclear flattening and 3D surface area (**Fig. 5E-G**). We detected cleaved pSTAT6 proteins in the nucleus, in line with previous reports ^47,48^.

**Figure 6.**
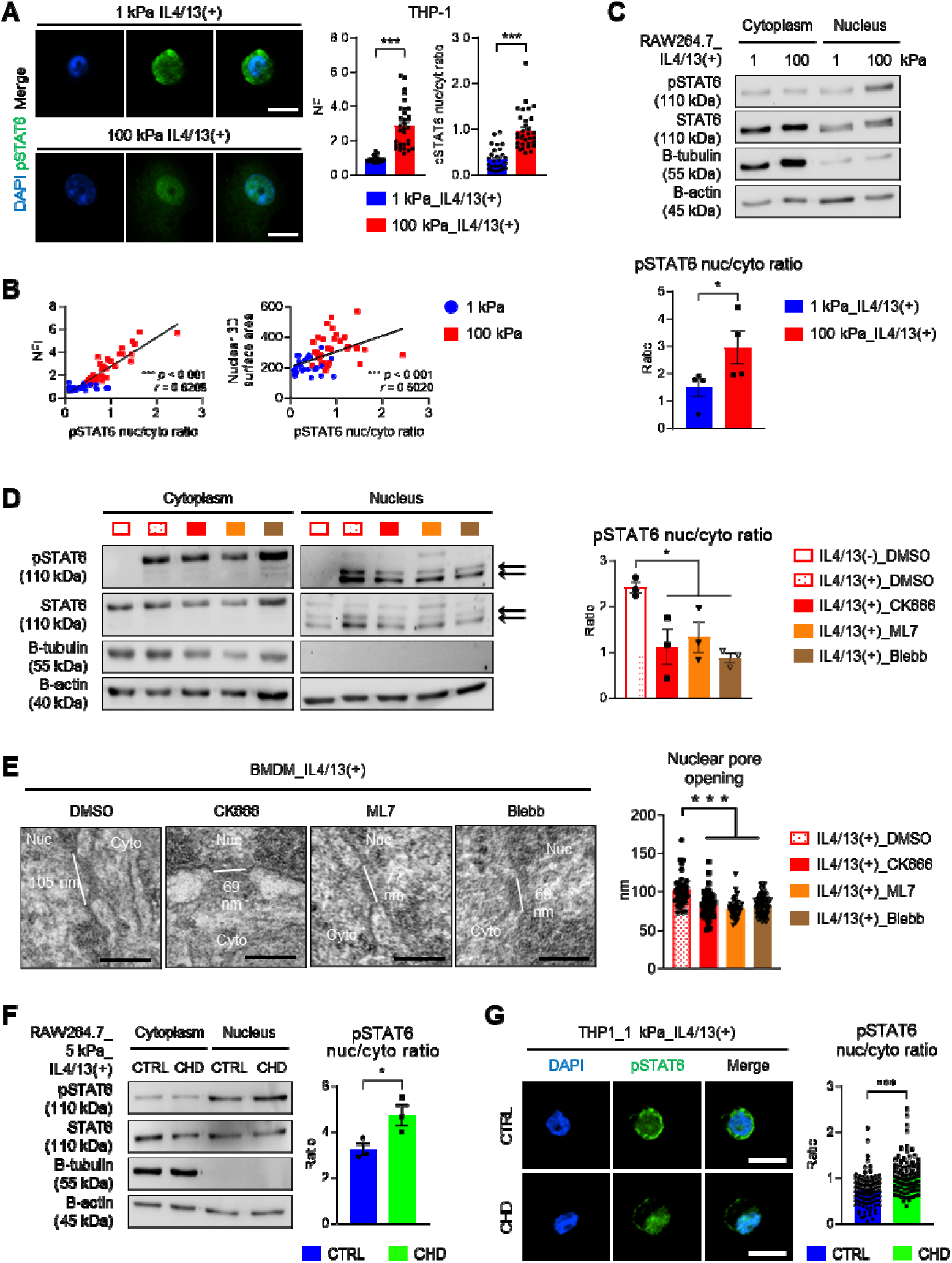
Nuclear pore opening and nuclear translocation of phosphorylated STAT6 increase via cytoskeletal-mediated nuclear tension increase. (**A**) Representative images of immunostaining of phosphorylated STAT6 (pSTAT6, green) and DAPI (blue) in THP-1 cells cultured on 1 kPa or 100 kPa PA gels in the presence of IL4/13 (scale bars = 10 µm). Nuclear flattening index (NFI) and nuclear (nuc)-to-cytoplasmic (cyto) ratio of pSTAT6 were measured and presented as mean ± SEM (*n* = 30 for 1 kPa_IL4/13(+) and *n* = 32 for 100 kPa_ IL4/13(+); \*\*\**p* < 0.001; Mann-Whitney test). **(B)** The correlations between nuclear-to-cytoplasmic ratio of pSTAT6 and NFI or nuclear 3D surface area were analyzed (one-tailed *p*-values and Spearman’s *r* are shown). **(C)** Representative Western blots and quantification of pSTAT6, STAT6, B-tubulin, and B-actin in cytoplasmic and nuclear fractions of RAW264.7 cells cultured on either 1 kPa or 100 kPa PA gel with IL4/13 treatment for 12 hours. The ratios of nuclear to cytoplasmic expressions of pSTAT6 normalized to B-actin are presented as mean ± SEM (*n* = 4 replicates; * *p* < 0.05; unpaired Student *t*-test). ns, not significant. **(D)** Representative Western blots and quantification of pSTAT6, STAT6, B-tubulin, and B-actin in cytoplasmic and nuclear fractions of BMDMs treated with DMSO, CK666 (200 µM), ML7 (25 µM), or Blebb (50 µM) in the absence or presence of IL4/13. Bands for cleaved pSTAT6 and STAT6 are indicated by arrows (approximately 85 to 75 kPa). The nuclear to cytoplasmic ratios of pSTAT6 expressions normalized to B-actin are presented as mean ± SEM (*n* = 3 replicates; \**p* < 0.05 and \*\**p* < 0.005; one-way ANOVA). ns, not significance. **(E)** Representative transmission electron microscopy (TEM) images of nuclear pores in BMDMs treated with DMSO, CK666, ML7, or Blebb in the presence of IL4/13 (scale bars = 100 nm). White lines indicate the lengths of opened nuclear pores. The mean ± SEM of nuclear pore sizes was calculated from 35 pores of 14 DMSO-treated cells, 47 pores of 11 CK666-treated cells, 38 pores of 10 ML7-treated cells, and 56 pores of 10 Blebb-treated cells (*** *p* < 0.001; Kruskal-Wallis test). **(F)** Representative Western blots and quantification of pSTAT6, STAT6, B-tubulin, and B-actin in cytoplasmic and nuclear factions of RAW264.7 cells cultured on 5 kPa PA gels pretreated without (CTRL, control) or with trans1-2 cyclohexanediol (CHD, 0.5%) for 1 h and then post-stimulated with IL4/13 for 1h. Normalized pSTAT6 expression data are presented as mean ± SEM (*n* = 3 replicates; * *p* < 0.05; unpaired Student *t*-test). **(G)** Representative images of immunostaining of pSTAT6 (green) and DAPI (blue) in THP-1 cells cultured on 1 kPa PA gels after treatment with CHD in the presence of IL4/13 (scale bars = 10 µm). Nuclear-to-cytoplasmic ratio (nuc/cyt) of pSTAT6 were measured and presented as mean ± SEM (*n* = 90 cells/group; \*\*\**p* < 0.001; Mann-Whitney test).

The interaction of TFs with the actin cytoskeleton or the direct coupling of the cytoskeleton to nuclear pore complexes (NPCs) that open in response to mechanical forces play a role in the nuclear translocation of TFs and transcriptional coactivators ^49–53^. Emerging studies have demonstrated that NPC is sensitive to nuclear deformation, and alterations in nuclear tension induced by mechanical stress or hypo-osmotic shock lead to immediate changes in NPC opening, influencing nuclear import of mechanosensitive TFs like YAP ^51,54^. To assess whether cytoskeletal-mediated nuclear deformations affect NPC opening, we measured NPC diameter in IL4/13-treated BMDMs before and after treatment with actomyosin inhibitors by transmission electron microscopy (TEM) (**Fig. 6E and Fig. S15**). Comparative analysis revealed that nuclear pores were significantly more closed following actomyosin inhibitor treatment, demonstrating that podosome-like actin dot structures and actomyosin-mediated force transmission led to nuclear deformation and subsequent NPC opening. To further substantiate that NPC opening facilitates the nuclear translocation of pSTAT6, we examined the subcellular localization of pSTAT6 through immunostaining and nuclear/cytoplasmic protein fractionation in THP-1 or RAW264.7 cells cultured on compliant substrates after treatment with trans1-2 cyclohexanediol (CHD), which disturbs the nuclear pore permeability through disrupting FG nucleoporins, known as a nuclear pore dilating agent ^51,55^. Even on compliant substrates (1 or 5 kPa), the nuclear import of pSTAT6 in IL4/13-treated macrophages was augmented when the nuclear pore barrier was disrupted (**Fig. 6F, G**). These findings indicate that the nuclear translocation of pSTAT6 and, consequently, M2 macrophage activation were promoted in macrophages cultured on rigid substrates due to NPC opening, which was caused by cytoskeletal-mediated nuclear tension increase and nuclear deformation.

### Chromatin remodeling and LADs reduction are linked to cytoskeletal-mediated nuclear mechanosensing

The modulation of heterochromatin condensation and positioning can occur through nuclear deformation induced by increased cytoskeletal process and actomyosin contractility, as reported in previous studies ^56–60^. In our exploration of the regulatory effects of cytoskeletal-mediated nuclear tension on the nuclear translocation of pSTAT6, we sought to determine whether chromatin remodeling is also influenced. We observed that expression levels of peripheral H3K9me2/3 and LADs were drastically reduced upon rigid substrates (**Fig. 2 and Fig. S6**). To examine whether the reduction in heterochromatin is specifically mediated by cytoskeletal processes, we assessed LADs in IL4/13-treated BMDMs cultured on 100 kPa PA gels following treatment with the inhibitors. Treatment with inhibitors led to increased peripheral enrichment of the heterochromatin mark, H3K9me2 (**Fig. 7A**), concomitant with a reduction in nuclear surface tension (**Fig. 5H-J**). Additionally, Dam-mediated m^6^A residues displayed a higher accumulation at the nuclear lamina of HEK-293 cells on 100 kPa PA gels after CK666 treatment (**Fig. 7B**). These findings suggest that cytoskeletal-mediated force transmission to the nucleus has a significant impact on chromatin remodeling.

**Figure 7.**
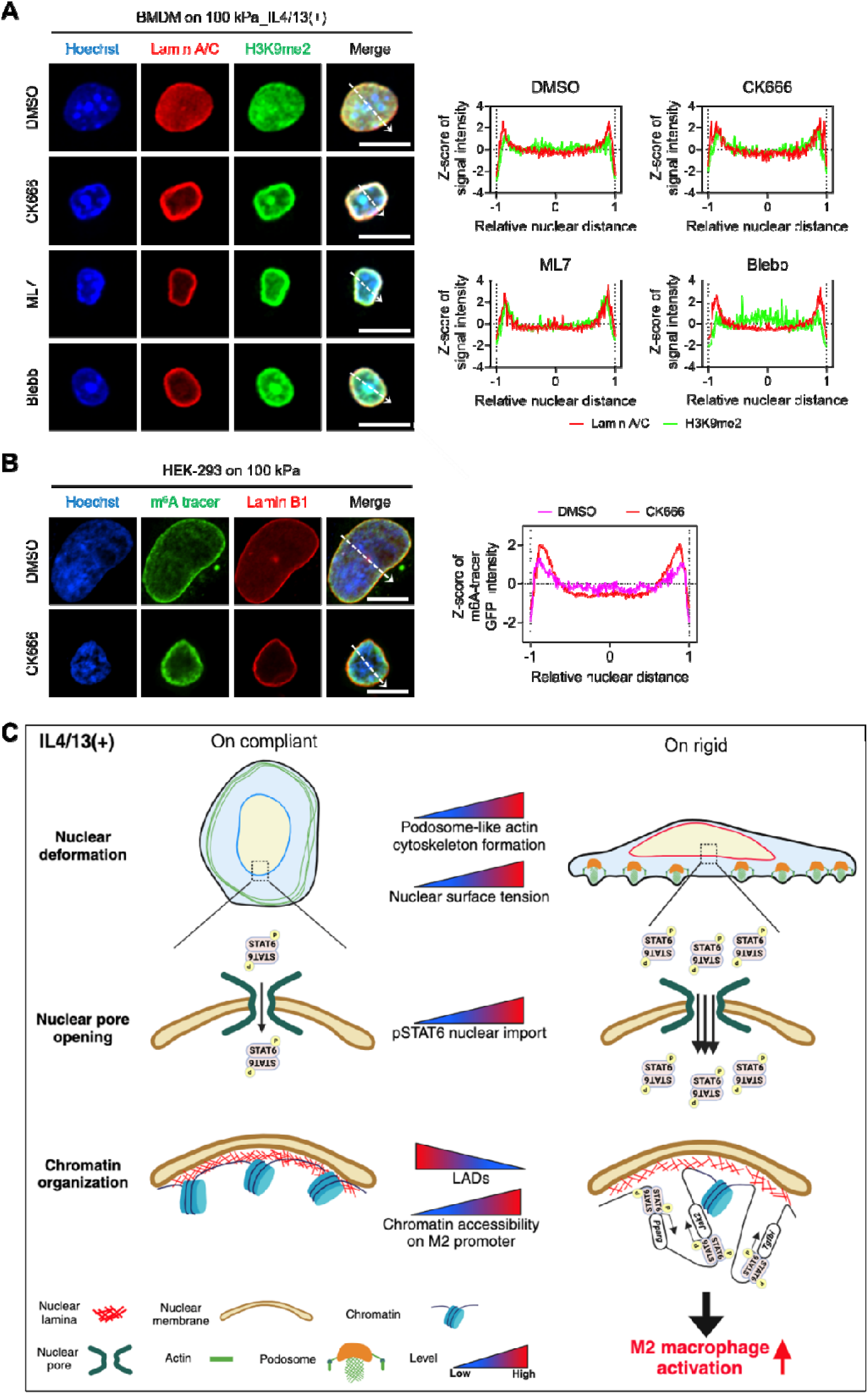
Chromatin remodeling is facilitated upon rigid matrices via cytoskeletal-mediated nuclear mechanosensing, driving M2 activation. (**A**) Representative co-immunostaining images of Lamin A/C (red), H3K9me2 (green), and Hoechst-33342 (blue) in BMDMs cultured on the 100 kPa_IL4/13(+) condition after treatment with CK666, ML7, blebbistatin (Blebb), or DMSO as a vehicle (scale bars = 10 µm). Dashed arrows indicate positions of the line signal intensity profiles. **(B)** Representative images of Hoechst-33342 (blue), m^6^A tracer (GFP, green), and Lamin B1 immunostaining (red) in HEK-293 cells cultured on 100 kPa PA gels and co-transfected with the DD-DamWT-LMNB1-IRES2-mCherry plasmid and m^6^A-Tracer-NES plasmid, followed by CK666 treatment. The mean Z-scores of normalized m^6^A tracer GFP intensity are presented across the relative nuclear distance, as indicated by the dashed arrows (*n* = 30 nuclei per each condition). **(C)** A model illustrating a potential mechanism by which M2 macrophage activation is promoted under higher matrix rigidity conditions through nuclear mechanosensing. When macrophages sense high matrix rigidity, podosome-like actin structures are constituted, leading to cytoskeletal-mediated force transmission to the nucleus, resulting in nuclear deformation. Subsequently, increased nuclear surface tension triggers nuclear pore opening, facilitating nuclear translocation of TFs, such as pSTAT6. Concurrently, LADs decrease, and chromatin is reorganized in response to actomyosin-mediated nuclear tension increase, increasing the accessibility of pSTAT6 on M2-associated gene promoter regions.

In summary, when macrophages are cultured on rigid substrates, the development of podosome-like F-actin structures facilitates actomyosin-mediated force transmission to the nucleus. Our observations indicate that M2 macrophage activation is potentiated under conditions of high matrix rigidity through two primary mechanisms. First, in response to increased nuclear tension on rigid substrates, NPC opening enhances the nuclear translocation of pSTAT6. Second, concurrently, the chromatin is decondensed at M2-associated genomic loci, rendering them more accessible to STAT6 and other transcriptional co-factors (as depicted in **Fig. 7C**).

### Clinical implications: Rigidity-primed M2 macrophages exhibit immunosuppression and accumulate in rigid tumor microenvironment

In the context of most tumors, infiltrated macrophages are commonly associated with the M2 phenotype, contributing to an immunosuppressive microenvironment conducive to tumor growth ^61^. Coincidentally, the extracellular matrix (ECM) rigidity is generally elevated in these tumor settings ^15^. To comprehensively understand the correlation between the abundance of M2 macrophages and increased ECM rigidity in tumors, we investigated whether M2-activated macrophages under high matrix rigidity exhibit immunosuppressive characteristics compared to their counterparts on compliant substrates. Conducting a cytokine array of conditioned media (CM) from IL4/13-treated BMDMs cultured on either 1 kPa or 100 kPa PA gels (**Fig. 8A**), our analysis revealed that BMDMs cultured on rigid substrates released higher levels of pro-tumoral cytokines, including tissue inhibitor of metalloproteinase 1 (TIMP-1), Progranulin, C-C motif chemokine ligand 7 (CCL7), and angiopoietin-related protein 3 (ANGPTL3), all implicated in cancer cell migration, proliferation, and angiogenesis (**Fig. 8B, C**). Furthermore, those cultured on rigid substrates released lower levels of pro-inflammatory cytokines involved in neutrophil/monocyte chemotaxis (**Fig. 8B, C**). Remarkably, the most strikingly downregulated cytokine was IFN-γ, a central player in anti-tumor immunity ^62^. To ascertain whether these macrophages on high matrix rigidity exert a functionally tumor-favorable effect against T cell proliferation and activation, we treated pre-stimulated primary human T cells with the CM from BMDMs for 3 days (**Fig. 8A**). In comparison to T cells exposed to the CM from BMDMs cultured on compliant substrates (33.6 ± 3.0%), T cells exposed to the CM from BMDMs cultured on rigid substrates exhibited significantly reduced proliferation (24.7 ± 2.4%) (**Fig. 8D**), indicating that BMDMs cultured on rigid substrates possess a more pronounced immunosuppressive capacity.

**Figure 8.**
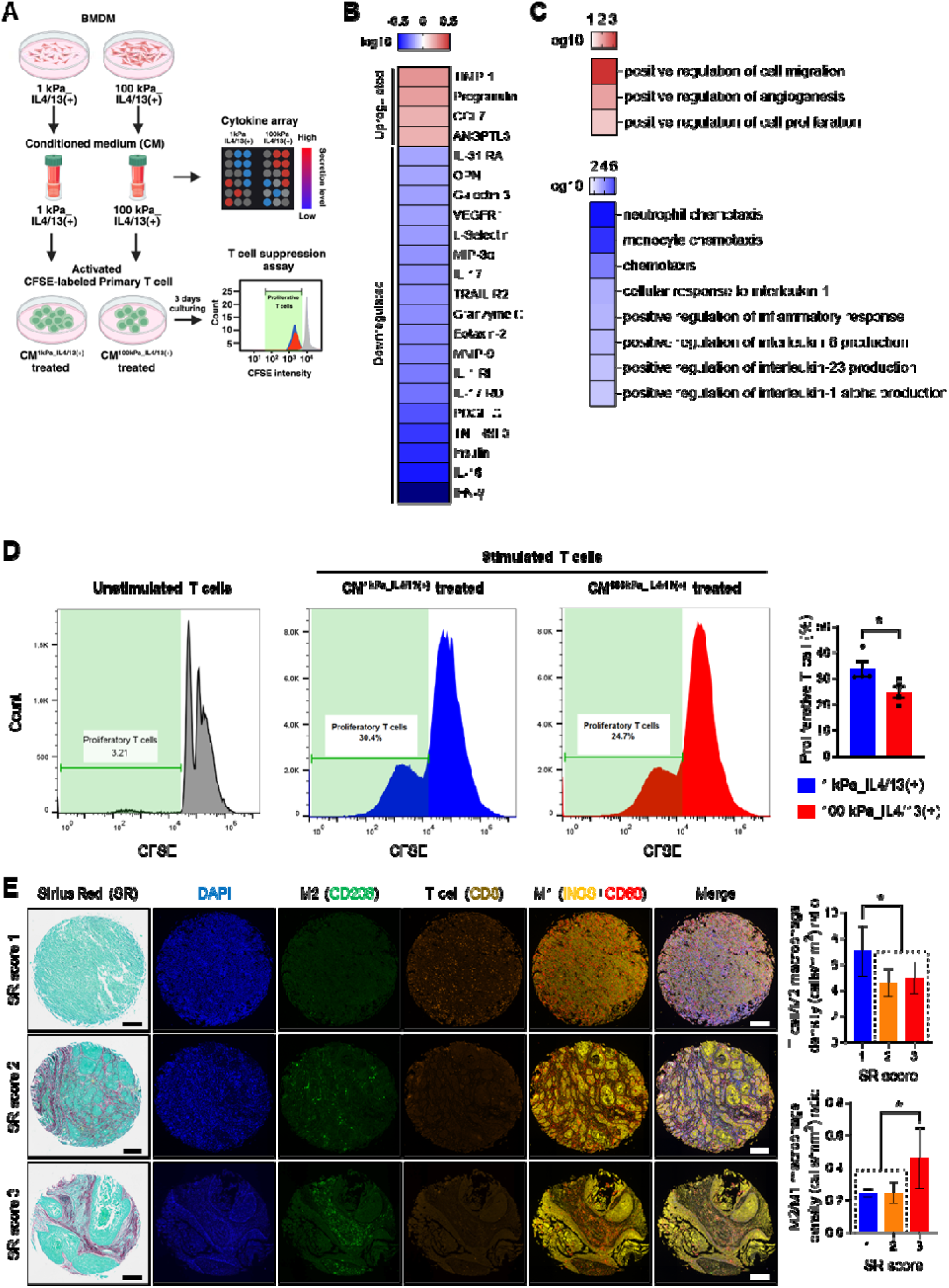
M2-activated macrophages exhibit enhanced immunosuppressive phenotypes under high environmental rigidity. (**A**) A schematic image illustrating the experimental design to evaluate the immunosuppressive characteristics of M2 macrophages under distinct substrate rigidity conditions. **(B)** Significantly increased (red) or decreased (blue) cytokines released from BMDMs cultured on 100 kPa PA gels, compared to that released from BMDMs cultured on 1 kPa condition (*p* < 0.05). The log10 normalized relative expression values are shown. **(C)** KEGG pathway analysis in which increased (red) or decreased (blue) cytokines are involved. The log10 normalized p-values are shown. **(D)** CFSE histograms detecting the inhibition of T cell proliferation after treatment with the conditioned medium (CM) of BMDMs cultured on 100 kPa, compared to that of BMDMs cultured on 1 kPa. Results are presented as mean ± SEM (*n* = 4 replicates; \**p* < 0.05; unpaired Student *t*-test). **(E)** Representative images of Sirius Red (SR) staining and multiplex immunofluorescence analysis for M2 macrophages (CD206; green), M1 macrophages (CD68; red, iNOS; yellow), cytotoxic T cells (CD8; orange), and DAPI (blue) in tissue microarrays of human head and neck squamous cell carcinoma tissues. Quantification graphs of the cell density (cells/mm^2^) ratio between cytotoxic T cells (CD8+) and M2 macrophages (CD206+) and between M2 and M1 macrophages (CD68+, iNOS+, CD206-) are shown. Results are presented as the mean ± SEM (1: *n* = 142, 2: *n* = 48, 3: *n* = 20; \**p* < 0.05; Mann-Whitney test or unpaired Student *t*-test, respectively).

To validate these findings in clinical samples, we assessed M1 and M2 macrophage and CD8^+^ T cell populations in patient tissues from head and neck squamous cell carcinoma. As tissue rigidity increased, inferred by collagen accumulation based on Sirius red staining, the ratio between M2 and M1 macrophage cell densities (M2/M1) significantly increased, while the ratio between cytotoxic T cells and M2 macrophages (T/M2) decreased (**Fig. 8E**). These results may suggest that M2 macrophage population was relatively increased as tissue rigidity increased, inhibiting anti-tumor T cell proliferation and M1 macrophage activation. Collectively, these findings propose that targeting matrix mechanics could potentially impede cancer progression by inhibiting M2 macrophage activation and ameliorating the immunosuppressive tumor microenvironment (TME).

## Discussion

Exploring the mechanisms by which high matrix rigidity potentiates M2 macrophage activation ^11,14^, this study revealed cytoskeletal-mediated increase in nuclear surface tension leads to nuclear deformation and pore opening, and subsequently chromatin reorganization. Our comprehensive analysis of ATAC-seq, evaluating genome-wide chromatin accessibility, demonstrated that both biochemical and mechanical stimuli cooperatively shape the epigenetic landscape for macrophage activation. Specifically, IL4/13-mediated JAK-STAT6 signaling and chromatin reorganization in STAT6-binding promoter regions of M2-associated genes were identified as key pathways for M2 macrophage activation under conditions of high matrix rigidity. Notably, despite some activation of IL4/13-mediated JAK-STAT6 signaling, nuclear import of pSTAT6 and accessibility of promoter loci decreased in macrophages cultured on compliant substrates compared to rigid ones. Thus, considering matrix mechanics is crucial for comprehensively understanding macrophage activation mechanisms and for modulating macrophage phenotypes for clinical applications.

This study revealed compelling evidence, supporting the notion that macrophages, when exposed to elevated substrate rigidity, induce the formation of podosome-like actin structures, which manifest a distinctive vinculin ring encircling the core F-actin, potentially through engagement with integrins. This phenomenon mirrors the observations in microvascular endothelial cells and human macrophages, responding to ECM rigidity ^63,64^. Given that the protrusion force of podosomes increases with rising matrix rigidity ^46,63^, our findings demonstrated that disrupting podosome formation (i.e., CK666 treatment) or actin traction force (i.e., ML7 and blebbistatin treatment) significantly inhibited nuclear mechanical alterations and M2 activation. This underscores the crucial role of podosome-like actin structures in the nuclear sensing of outside-in mechanical stresses. Moreover, this study, for the first time, explains how nuclear deformation leads to gene expression alterations via epigenetic regulations in macrophages by comprehensively analyzing the H3K9me2/3-marked heterochromatin or LADs distribution and genome-wide chromatin accessibility changes with varying matrix rigidity in the absence or presence of IL4/13.

Several limitations exist in this study. The full understanding of how podosome-like actin structures can transduce mechanical stresses into the nucleus and how alterations in nuclear mechanics modulate macrophage activation remain elusive. While it is known that the linker of nucleoskeleton and cytoskeleton (LINC) complex, composed of SUN-domain proteins and KASH-domain proteins across the inner and outer nuclear membranes, is implicated in linking the nuclear lamina/chromatin and cytoskeleton ^50,51^, the physical connection between podosome and the nucleus remains unknown ^65^. Our speculation posits that higher matrix rigidity increases cellular spreading through the augmentation of podosome-like actin structure formation, acting as tent nails. The more spreading cellular morphology decreases the total cellular height, dorsal cellular membrane compresses the nuclear membrane, and this compressive force leads to nuclear deformation, subsequently affecting the nuclear import of pSTAT6. Given that microtubules, another cytoskeletal component, regulate podosome formation ^44^ and stabilize podosome structures by passing through podosomes at the layer of myosin IIA in macrophages ^66^, the interplay between podosomes and microtubules, which have a direct connection with the LINC complex through Nesprins ^67^, is also likely involved in nuclear mechanosensing.

External forces transmitted to the nuclear membrane can influence chromatin positioning, thereby affecting gene expression ^25,68^. For instance, culturing cardiomyocytes on higher matrix rigidity reveals decreased peripheral enrichment of H3K9me3-marked heterochromatin in the nuclei ^56^. During confined migration, the nuclei in cancer cells and fibroblasts exhibit increased heterochromatin levels marked by H3K9me3 and H3K27me3 ^69^. Moreover, exposure to short-term uniaxial stretch induces a substantial reduction of H3K9me2/3 in epidermal progenitor cells, with the decreased heterochromatin levels returning to a steady state in the case of long-term stretch ^70^. In our study, changes were observed in the sequestering of H3K9me2/3-marked heterochromatin to the nuclear periphery, termed as LADs, and in the H3K9me2/3 expression levels dependent on matrix rigidity. Additionally, nuclear mechanosensing influences chromatin accessibility. Previous studies have reported that the nucleus of tumor cells is highly mechanosensitive to matrix rigidity, affecting the chromatin landscape and leading to a more tumorigenic phenotype in rigid matrices ^57,58^. Chromatin accessibility of fibroblasts and mesenchymal stem cells can also be reconstructed to attain distinct cellular phenotypes in response to mechanical stress such as matrix rigidity ^59^ and cyclic stretch ^60^. Our results suggest that both mechanical forces triggering nuclear mechanosensing and biochemical signals (i.e., IL4/13) cooperatively increase chromatin accessibility on promoter regions of genes associated with M2 macrophage activation, along with gene transcription levels. Interestingly, chromatin accessibility on promoter regions of M1-related genes was significantly decreased under the M2-potentiating condition, suggesting that chromatin organization and accessibility are exquisitely controlled in a context-dependent manner. How chromatin accessibility can be precisely regulated by complicated biochemical and mechanical stimuli remains open for further study, utilizing advanced techniques to capture genome conformation and chromatin interactions.

Not only matrix rigidity but also surface topography and geometry can influence M2 macrophage activation ^71^. Macrophages cultured on an array of microgrooves of varied dimensions tend to express more *Arg1* with an increased degree of elongation ^72^. Biomimetic wrinkle substrates that cause macrophage elongation promote *Arg1* expression and IL10 secretion *in vitro* ^73^. In 3D, fibrous geometries generated by electrospinning pores appear to alter macrophages, with a more pronounced M2 macrophage response to microscale fibers compared to nanoscale fibers ^74^. All these findings demonstrate that macrophages are highly mechanosensitive cells adopting their phenotypes depending on the environmental conditions. In the context of cancer, matrix rigidity enhances interactions between cancer cells and macrophages and M2-like macrophage accumulation in the TME ^16,75^. Treatment with matrix softening agents, such as β-aminopropionitrile, a lysyl oxidase inhibitor, changed the M1 to M2 macrophage ratio, increasing M1 macrophages while reducing the M2 macrophage population ^16^. Our findings suggest that M2-activated macrophages under high matrix rigidity exhibit a more pronounced T cell suppressive effect and secret higher levels of pro-tumorous cytokines, implicating that manipulating matrix rigidity may offer therapeutic opportunities against cancers. Furthermore, if there is a common mechanotransducive mechanism that potentiates M2 macrophage activation due to environmental mechanical stimuli, it may be possible to develop a therapeutic agent that induces the polarization of M2 macrophages towards M1 phenotypes, establishing an anti-immunosuppressive TME.

In summary, our work sheds light on the intricate interplay between matrix rigidity and nuclear mechanics during IL4/13-induced M2 macrophage activation and defines podosome-like actin-mediated force transmission as required for nuclear mechanosensing, leading to chromatin remodeling and increasing TF availability that induce the enhancement of M2 macrophage activation.

## Materials and Methods

### Fabrication of polyacrylamide (PA) gels

The preparation of PA gels followed the methodology outlined in a previous study ^76^. In brief, a mixture of 40% acrylamide solution (BIORAD, #1610140), 2% Bis-Acrylamide solution (BIORAD, #1610142), and distilled water was blended and adjusted to manufacture PA gels with varying rigidity. This PA gel solution was degassed for 15 minutes (min) to expedite polymerization and enhance gel uniformity. Subsequently, the gel solution was mixed with 10% ammonium persulfate (Sigma-Aldrich, #A3678) and TEMED (Sigma-Aldrich, #T9281), and promptly polymerized onto coverslip glass. To immobilize extracellular matrix proteins onto the surface of the PA gel, a hetero-bifunctional crosslinker, 0.5 mg/mL Sulfo-SANPAH (Sigma-Aldrich, #80332), was applied to the gels, followed by exposure to UV light. The gels were then coated with 0.05 mg/mL fibronectin (Sigma-Aldrich, #F0895) and incubated at 37 °C overnight to facilitate crosslinking.

### Isolation and primary culture of mouse bone marrow-derived macrophages (BMDMs)

Primary mouse BMDMs were isolated from the femurs of 5– to 8-week-old ICR mice following a previously reported protocol ^77^. The bone marrow cells were collected by flushing the femurs using serum-free alpha MEM media (WELGENE, #LM008-53). The removal of red blood cells was accomplished through lysis using eBioscience 1× RBC Lysis Buffer (Invitrogen, #00-4333-57). Following centrifugation and resuspension in alpha MEM media supplemented with 10% fetal bovine serum (FBS; CORNING, #35-015-CV) and 1% Penicillin-Streptomycin (Gibco, #15140163), the cells were filtered through a 40 µm cell strainer and incubated in the aforementioned media for 8 hours (h). Subsequently, non-adherent cells were transferred to cell culture dishes and cultured in alpha MEM containing 10% FBS, 1% Penicillin-streptomycin, and 30 ng/mL MCSF (Peprotech, #315-02) for 3 days to induce their differentiation into the macrophage phenotype.

For subsequent experiments, BMDMs were seeded onto PA gels and allowed to adhere for 12 h. Then, either 20 ng/mL IL4/13 (IL4: Peprotech, #214-14; IL13: Peperotech, #200-13) or 100 ng/mL IL10 (Peprotech, #210-10) was added for a 12-h period to induce M2 activation. Following activation, the cells were harvested and utilized for further investigations. In certain experiments, pretreatment with CK666 (TOCRIS, #3950), ML7 (TOCRIS, #4310), and Blebbistatin (TOCRIS, #1760) was performed 1 h prior to IL4/13 stimulation. JIB-04 (TOCRIS, #4972) was administered as a pretreatment for 12 h before IL4/13 induction.

### Cell lines and culture

RAW264.7 and HEK-293 cells were maintained in DMEM high-glucose media (WELGENE, #LM001-05) supplemented with 10% FBS, and 1% of Penicillin-Streptomycin. THP-1 cells were cultured in RPMI1640 (WELGENE, #LM011-03) with 10% FBS and 1% of Penicillin-streptomycin. To induce macrophage differentiation in THP-1 cells, 100 ng/mL Phorbol 12-myristate 13-acetate (PMA; BioGems, #1652981) was added, and the cells were cultured for 1 day. Following differentiation, THP-1 cells were subjected to serum and PMA deprivation before undergoing M2 activation induction.

### QuantSeq 3’mRNA sequencing

RNA samples were extracted from BMDMs using the Ribospin RNA extraction kit (GeneAll, #304-150). To eliminate potential genomic DNA contamination, the RNA samples were treated with DNase I, RNase-free (New England Biolabs, #M0303S). Following quality control assessment, all RNA samples meeting the criteria were subjected to library preparation, and QuantSeq 3’mRNA-seq was conducted by ebiogen Inc. (South Korea). A total of approximately 19.5 million reads, corresponding to 23,282 unique genes, were sequenced, trimmed, and aligned to the mouse reference genome (GRm38/mm10). The acquired processed data was analyzed employing ExDEGA software (v4.0.3, ebiogen, South Korea). For downstream analyses, we exclusively considered 2,866 genes with a normalized log2 expression greater than 4 in all conditions. Differentially expressed genes (DEGs) were identified if the fold change between groups exceeded 1.5. Enrichment analysis of Gene Ontology (GO) terms was carried out using DAVID (https://david.ncifcrf.gov/) ^78^.

### Assay for transposase-accessible chromatin using sequencing (ATAC-seq)

For ATAC-seq, sample preparation followed previously established methods ^79^. Briefly, BMDMs were detached from PA gels using the Macrophage Detachment Solution (PromoCell, #C-41330). Subsequently, cell count was determined using a hematocytometer, and 50,000 cells were lysed using lysis buffer consisting of 10 mM TrisHCl pH7.4, 10 mM NaCl, 3 mM MgCl_2_, and 0.1% IGEPAL CA-630 (Sigma-Aldrich, #18896). The lysate was centrifuged, and supernatants were removed. The resulting cell nuclei were subjected to incubation for 30 min with Tagment DNA TDE1 Enzyme and Buffer (Illumina, #FC-121-1030). This step fragmented the chromatin and inserted sequencing adapters. Following the reaction, the transposed DNA was purified using the MinElute PCR Purification Kit (Qiagen, #28004). ATAC-seq was carried out by Macrogen Inc. (South Korea). On average, 40,234,171 reads per sample were sequenced and aligned to the GRm38/mm10 mouse reference genome. Subsequent analysis of processed peak data utilized ExDEGA software (v4.0.3, ebiogen, South Korea). For downstream analyses, exclusive focus was placed on 2,203 regions exhibiting a normalized log2 peak value exceeding 1 across all conditions. Distinct regulatory elements (DREs) were identified when the fold change between groups surpassed 1.5. GO enrichment analysis was performed using the DAVID platform. Discovery of de novo transcription factor (TF) binding motifs within ATAC-seq-defined regions was carried out using the HOMER software ^80^. Enrichment of TF binding motifs, including MA0520.1 (STAT6) and MA0490.2 (JunB), in open chromatin regions aligned to the mouse reference genome (GRm38/mm10) was assessed by calculating log2 normalized coverage levels of peak area using TFmotifView at default options ^81^. The Integrative Genome Viewer (IGV, http://igv.org/) was employed to visualize the genome track corresponding to the gene locus.

### Computational modeling of nuclear tension

We employed a structural mechanics module-based modeling and simulation to assess the surface tension of deformed nuclei in response to changes in matrix rigidity. The finite element method (FEM) implemented within the COMSOL Multiphysics software was used for this purpose. The computational framework was established in a 2D geometry with a symmetric axis along the z-axis, later extrapolated to create a 3D structural model using a revolved geometry approach. The mesh of the model was generated using free triangular elements with a maximum element growth rate set at 1.3. This resulted in a total of 7,238 elements, with an average element quality of 0.9416, as indicated by mesh statistics. In our simulation, we modeled a cell adhering to either a compliant or rigid substrate as a hemisphere with a radius of 12 µm. Inside the cell, the nucleus was represented as an ellipsoidal shape with a vertical radius of 4 µm and a horizontal radius of 3.5 µm. To characterize the mechanical properties of the cell, we employed a linear elastic material with a viscoelastic component, specifically a generalized Maxwell model, summarizing its mechanical characteristics in **Table S1**.

We simulated nuclear deformation by applying pulling forces to the bottom of the cell that adheres to the deformable substrates. This resulted in the generation of pressure by the cell membrane, which was assumed to be connected via the intracellular cytoskeletons. We compared the forces applied to the surface of deformed nuclei under different cell adhesion conditions. The analysis of deformation was conducted using a stationary model to simplify and address steady-state problems. The external force, mimicking focal adhesion, was simulated by a combination of prescribed displacements and boundary loads applied to the cell. The direction of cell strain was assumed to be outward due to the increase in matrix rigidity ^82,83^. Prescribed displacement points were strategically located at various distances along the cell bottom (12, 8, and 4 µm from the center of the cell), with displacement ratios of 1, 0.6, and 0.3 applied to both substrates. Additionally, a coupled body load, directly proportional to the prescribed displacement (scaled by a factor of 10-3 N/m), was applied to the cell body. The bottom of the cell was constrained as a boundary condition, ensuring zero displacement in the perpendicular direction.

### Cytokine array

The conditioned medium was collected from BMDMs cultured on 1 kPa or 100 kPa substrates after pre-treatment with IL4/13 for 12 h. Subsequently, the collected medium was centrifuged at 2000 rpm for 3 min at 4 °C. The resulting supernatant was used for the cytokine array analysis, employing the Mouse L308 Array, Membrane (RayBiotech, #AAM-BLM-1-4). The analysis and data acquisition were conducted by ebiogen Inc. (South Korea).

### Quantitative reverse transcription polymerase chain reaction (qRT-PCR)

Total RNA was extracted using the Ribospin RNA extraction kit (GeneAll, #304-150). 500 ng of total RNA was subjected to reverse transcription using the iScript cDNA Synthesis Kit (BIORAD, #1708891). For amplicon detection, the SensiMix SYBR Hi-ROX Kit (meridianBIOSCIENCE, #QT605-05) was used, with gene expression normalized against a house keeping gene GAPDH. The qRT-PCR reactions were performed in triplicate on a StepOnePlus Real-Time PCR System (Applied Biosystems, #4376592). The cycling conditions consisted of an initial denaturation step at 95 °C for 10 min, followed by 40 cycles of denaturation at 95 °C for 15 seconds (s), annealing at 55 °C for 15 s, and extension at 72 °C for 30 s. The quantification of mRNA transcription levels was achieved using the ddCt method, with values expressed relative to the mean dCt of control samples. Primer sequences are provided in **Table S2**.

### Western blot

The cellular lysates were prepared using Protein Extraction Solution (ELPISBIO, #EBA-1149) supplemented with 1% Halt Protease and Phosphatase Inhibitor (ThermoFisher, #78442), followed by a 30-min incubation on ice. The supernatant containing proteins was collected after centrifugation. For nuclear and cytoplasmic fractionation, NE-PER Nuclear and Cytoplasmic Extraction Reagents (ThermoFisher, #78835) was utilized. Protein concentration was determined using Pierce BCA Protein Assay Kit (ThermoFisher, #23225). Subsequently, an equivalent amount of protein for each sample was mixed with Laemmli’s 5× Sample Buffer (ELPISBIO, #EBA-1052) and heated at 95 °C for 5 min. The mixture was then subjected to SDS-PAGE electrophoresis. Afterwards, proteins were transferred onto either a Nitrocellulose Membrane 0.45 µm (BIORAD, #1620115) or a 0.2 µm membrane (BIORAD, #1620112) for small-sized proteins (<20 kDa). Overnight incubation at 4 °C was conducted for the first antibody reactions. Subsequently, membranes were probed with HRP-conjugated anti-mouse (Cell Signaling, #7074) or anti-rabbit (Cell Signaling, #7076) secondary antibodies for 1 h. Detection was achieved using the Chemiluminescent Substrate Kit (ThermoFisher, #34580). Detailed information regarding antibodies was summarized in **Table S3**.

### Cellular morphometric analysis

For visualization of the cell membrane and nucleus, CellMask Plasma Membrane Stains (Invitrogen, #C10046) were used to label the cell membrane, while DAPI (Sigma-Aldrich, #D9542) or Hoechst-33342 (Invitrogen, #H3750) were employed for nuclear staining. Morphometric analysis of both nuclear and cellular 2D areas was executed using the NIH ImageJ software. To calculate the density of actin dots (actin dots/100um^2^), the cellular surface area was measured, along with the quantification of the number of actin dots per cell, utilizing an ImageJ macro previously reported ^84^. To create a 3D nuclear dataset, cells stained with DAPI or Hoechst-33342 were captured using Z-stack imaging. Subsequently, calculations for nuclear 3D surface area were performed using the NIS-Elements Imaging Software (Nikon, Japan). Nuclear flattening was determined based on a previously established method ^79^. This metric was defined as the ratio a/h, where ‘a’ represents the longest diameter of the 2D nucleus counterstained with DAPI or Hoechst-33342, and ‘h’ signifies the nuclear height extracted from the Z-stack 3D image.

### Immunofluorescence

For the immunofluorescence analysis of Lamin A/C, H3K9me2, and H3K9me2/3, the cells were fixed using 100% methanol for 5 min at –20 °C. In contrast, for the immunostaining of Vinculin, and phosphorylated STAT6 (pSTAT6), the cells were fixed with 4% paraformaldehyde (PFA) in phosphate-buffered saline (PBS) for 15 min at room temperature, followed by permeabilization using 0.1% Triton X-100 in PBS for 10 min. Afterwards, the cells were blocked with 1% BOVINE SERUM ALBUMIN (BSA) FRACTION V (GeneAll, #SM-BOV-100) in PBS containing 0.1% Triton X-100 for 60 min. Following the overnight incubation with the primary antibody at 4 °C, secondary antibodies specific to rabbit (JacksonImmunoResearch, #711-025-512) and mouse (JacksonImmunoResearch, #715-025-150) were introduced for 1 h at room temperature. Nuclei were counterstained using DAPI or Hoechst-33342, while actin was labelled with Alexa Fluor 488 phalloidin (Invitrogen, #A12379). Detailed antibody information was summarized in **Table S4**.

To determine the nuclear to cytoplasmic ratio of pSTAT6, the mean fluorescence intensity in the nucleus was measured and divided by the mean fluorescence intensity in the cytoplasm. To assess the peripheral enrichment of Lamin A/C, H3K9me2, and H3K9me2/3, a line was drawn along the long axis of nuclei. The signal intensity was then quantified using ImageJ, and the actual nucleus distance was transformed into a normalized relative distance scale. In this scale, the center of the nucleus was assigned 0, while the periphery of the nucleus was designated –1 and 1. This normalized relative distance was divided into 100 segments, and the signal intensity within each segment was averaged to generate a graph. A total of 30 measurements were performed for these analyses.

### DNA adenine methyltransferase identification (DamID)

The DD-DamWT-LMNB1-IRES2-mCherry plasmid (Dam-LMNB1; addgene, #159601) and m^6^A-Tracer-NES plasmid (m6A-Tracer; addgene, #159607) were simultaneously transfected into HEK-293 cells using Lipofectamine 3000 Transfection Reagent (Invitrogen, #L3000008). After 48-h co-transfection, the cells were re-seeded onto 1 or 100 kPa PA gels. After the cells were attached on the gels, 0.5 µM Shield-1 (MedChemExpress, #HY-112210) were treated for 12 h. The fluorescence signal was visualized using confocal microscopy.

### Transmission electron microscopy (TEM)

2 × 10^6^ BMDMs were treated with either DMSO or inhibitors (CK666, ML7, Blebbistatin) in the presence of IL4/13. The cells were fixed using a solution of 2% glutaraldehyde and 2% PFA in PBS for 1 h at room temperature. Following fixation, the cells were gently scraped and subsequently centrifuged to form cell pellets. These pellets were then resuspended in the fixation solution and incubated for 12 h at 4 °C. Subsequent steps involved dehydration and infiltration with propylene oxide. The samples were embedded using the Poly/Bed 812 Embedding Kit (Polysciences, #08792-1) and sectioned to a thickness of 200 to 250 nm. Initial observations were conducted on sections stained with toluidine blue (Sigma-Aldrich, #T0394) using a light microscope. For detailed analysis, 80 nm sections were subjected to double staining with uranyl acetate (Electron Microscopy Sciences, #541-09-03) for 20 min and lead citrate (Fisher scientific, #6107-83-1) for 10 min. The stained sections were then cut using a diamond knife, affixed to a Leica EM UC7 (Leica MICROSYSTEMS, Austria), transferred onto copper grids, and finally examined using a TEM HT7800 (HITACHI, Japan) operating at 100 kV. ImageJ software was employed for measurement of nuclear pore opening in BMDMs (n = 2 per group).

### Modulation of nuclear pore permeability

Cells were seeded on fibronectin-coated PA gels with a rigidity of 1 or 5 kPa for 12 hrs. To increase nuclear pore permeability, 0.5% trans-1,2-cyclohexanediol (CHD) (Sigma-Aldrich, #141712) was applied for 1 h prior to IL4/13 stimulation for 1 h. Subsequently, THP-1 cells were subjected to immunofluorescent staining for pSTAT6 and Hoechst-33342, and RAW264.7 cells were used for Western blots after proceeding nuclear cytoplasmic fraction. The nuclear/cytoplasmic pSTAT6 ratio was quantified using the methodology described above.

### *In vitro* efferocytosis assay

The efferocytosis assay was conducted following methodologies outlined in prior studies ^85–87^. Jurkat cells were exposed to UV irradiation (80 uW/mn^2^) for 15 min, followed by an incubation at 37 °C for 2 to 3 h to induce apoptosis. Using flow cytometry, we evaluated that this methodology consistently produced cell populations in which approximately 90% of cells exhibited positive staining with annexin V, while less than 5% displayed positive staining for propidium iodide (PI). Apoptotic Jurkat cells (ACs) were then labeled with the tracking dye PKH67 (Sigma-Aldrich, #MIDI67), and any excess dye was removed thorough two washes with 1% BSA in HBSS (WELGENE, #L003-02). The labeled ACs were added to BMDMs plated on cell culture dishes at a 1:5 ratio of BMDMs to ACs, and the co-culture was incubated for 45 min at 37 °C. Subsequently, after a PBS wash to eliminate any surplus ACs, the BMDMs were fixed using 1% PFA for 10 min. Following fixation, the cells were stained with Alexa Fluor 546 phalloidin (Invitrogen, #A22283) and DAPI to visualize F-actin and nuclei, respectively. The assessment of efferocytotic capacity was performed by determining the percentage of PKH67-positive BMDMs relative to the total BMDM population. For qRT-PCR analysis, 1 h after adding ACs to the BMDMs, the BMDMs were rinsed with condition medium and then cultured for an additional 6 h, followed by RNA extraction for subsequent analysis.

### T cell suppression assay

The primary T cells were obtained from peripheral venous blood of a healthy human donor using the EasySep™ Direct Human T cell Isolation Kit (STEMCELL TECHNOLOGIES, #19661) following the manufacturer’s instructions. These isolated cells were then labeled with the CellTrace™ CFSE Cell Proliferation Kit for flow cytometry (Invitrogen, #C34554) and primarily cultured in CTS AIM V SFM (Gibco, #0870112DK) supplemented with 10 ng/mL of Human Recombinant IL-2 (STEMCELL TECHNOLOGIES, #78220) and ImmunoCult™ Human CD3/CD28 T Cell Activator (STEMCELL TECHNOLOGIES, #10971) for a duration of 3 days. Conditioned medium obtained from BMDMs seeded on 1 kPa and 100 kPa PA gels, which had been treated with IL4/13 for 12 h, was centrifuged. The resulting supernatant was concentrated using Amicon Ultra-15 Centrifugal Filter Unit (Millipore, #UFC901024). Subsequently, the primary T cells were re-cultured using this concentrated medium. After 3 days of re-culture, the level of CFSE in proliferating T cells was assessed using the BD Accuri™ C6 Plus Personal Flow Cytometer (BD Biosciences, USA).

### Multiplex immunofluorescence and collagen deposition assessment in tissue microarrays

Tissue microarrays (TMAs) comprising 210 head and neck squamous cell carcinoma samples (OR208a, OR601c) were procured from TissuArray.Com (USA). Sirius Red staining was performed on all TMAs, and images were acquired using a slide scanner (OlyVIA 3.2, VS2000 ASW, Olympus). H&E-stained images were obtained from TissueArray.Com. Quantitative evaluation of Sirius Red staining was conducted as previously described with minor modifications ^88^. The proportion of Sirius Red-positive area was calculated in the tumor-surrounding stroma and scored as follows: 1, 0%-25%; 2, 26%-50%; 3, 51%-100%. Multiplex immunofluorescence targeting M2 macrophages (CD206+), M1 macrophages (CD68+, iNOS+, CD206-), cytotoxic T cells (CD8+), and DAPI, along with the quantification of each cell densities (cells/mm^2^), was performed by BioActs Co. (Incheon, South Korea) and data were provided.

### Statistical analysis

The bar graphs illustrating experimental results were presented as mean ± standard error of the mean (SEM). Statistical analyses were conducted using Prism 9 software (v9.4.0, GraphPad, USA). The Shapiro-Wilk test was employed to assess the normality of the data. For datasets exhibiting a normal distribution, statistical significance was assessed using the student’s *t*-test for two-group comparisons or ordinary one-way analysis of variance (ANOVA) for multiple group comparisons. In cases where the data did not conform to a normal distribution, non-parametric tests, such as the Mann-Whitney test for two-group comparisons and the Kruskal-Wallis test for multi-group comparisons, were utilized. The levels of significance were denoted as follows: * for *p* < 0.05, ** for *p* < 0.005, and *** for *p* < 0.001, indicating statistically significant results.

### Author contribution

SJS performed experiments, analyzed and interpreted the data, and drafted the manuscript; BB, KT, MY, and YJK performed experiments; SHK analyzed the sequencing data; DK conducted computational modeling of nuclear mechanics; DHK conceptualized the study, conducted computational modeling of nuclear mechanics, and edited the manuscript; JHL conceptualized the study, designed the experiments, secured funding for the research and co-supervised the study; JH conceptualized the study, designed the experiments, analyzed and interpreted the data, wrote and edited the manuscript, secured funding for the research and co-supervised the study; HWK conceptualized the study, designed the experiments, edited the manuscript, secured funding for the research and supervised the study.

## Acknowledgement

This work was supported by grants from the National Research Foundation of Korea (2021R1C1C1003904 to JH, 2021R1A5A2022318 to HWK, and 2V09840-23-P024 to DK).

