## supplementary figs 1-15, tables 1-4 for "Matrix-rigidity cooperates with biochemical cues in M2 macrophage activation through increased nuclear deformation and chromatin accessibility"

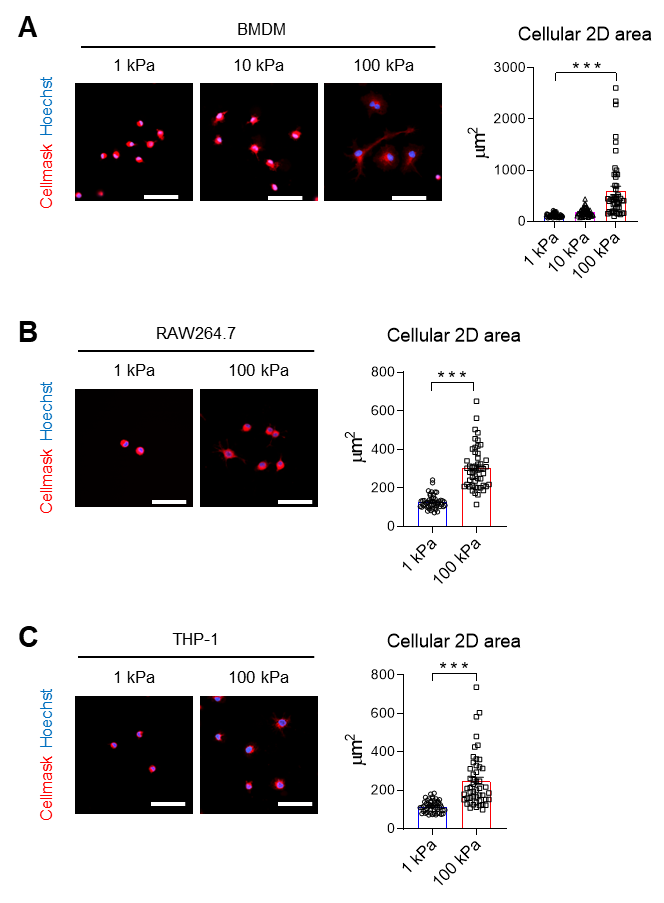

**Supplemental Figure 1. Macrophages undergo morphological changes in response to varying matrix stiffness.** **(A-C)** Representative images of CellMask (red) and Hoechst-33342 (blue)-stained BMDMs **(A)**, RAW264.7 **(B)**, and THP-1 cells **(C)** cultured on PA gels with varying matrix rigidity (1, 10, or 100 kPa). Scale bars represent a length of 50 µm. Quantification of cellular 2D projection area is presented as mean ± SEM (*n* = 30 cells/group; ****p* < 0.001; Kruskal-Wallis test or Mann-Whitney test).

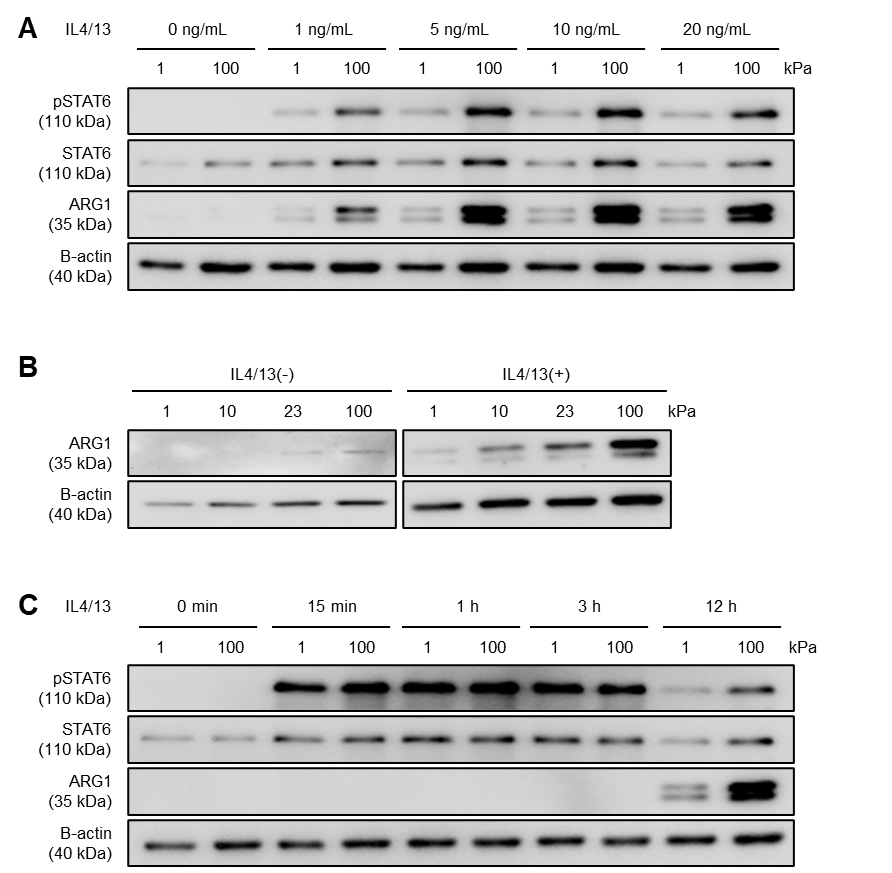

**Supplemental Figure 2. IL4/13-mediated induction of M2 macrophage markers is enhanced by elevated matrix rigidity. (A)** Western blot images showing pSTAT6, STAT6, ARG1, and B-actin in BMDMs exposed to varying concentrations of IL4/13 (0, 1, 5, 10, and 20 ng/mL) for 12 hours on either 1 kPa or 100 kPa PA gels. **(B)** Western blots of ARG1 and B-actin in BMDMs cultured on substrates with different rigidity levels (1, 10, 23, and 100 kPa) with or without IL4/13 treatment for 12 hours. **(C)** BMDMs were treated with IL4/13 (20 ng/mL) for 0, 15 minutes, 1 hour, 3 hours, or 12 hours on either 1 kPa or 100 kPa PA gels. Western blot images of pSTAT6, STAT6, ARG1, and B-actin are shown.

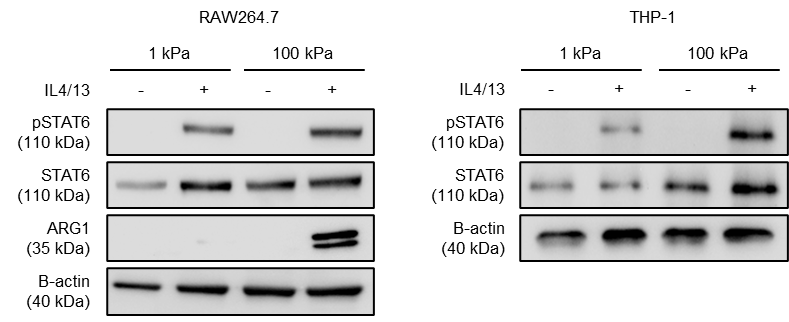

**Supplemental Figure 3. IL4/13-induced upregulation of M2 macrophage markers is enhanced in high rigidity matrices.** Representative Western blot images of pSTAT6, STAT6, ARG1, and B-actin in RAW264.7 and THP-1 cells cultured on PA gels with rigidity levels of 1 kPa or 100 kPa, with or without IL4/13 treatment for a duration of 12 hours.

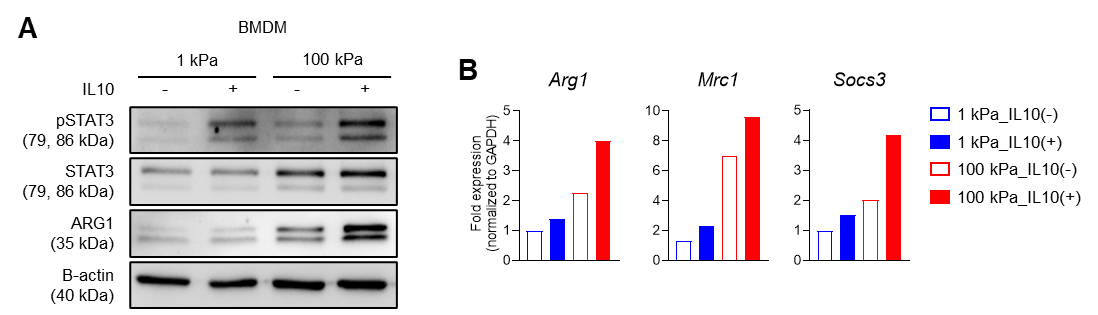

**Supplementary Figure 4. IL10-driven induction of M2 macrophage markers is augmented in response to increased matrix rigidity. (A)** Representative Western blot images of pSTAT3, STAT3, ARG1, and B-actin in BMDMs cultured on 1 kPa or 100 kPa PA gels, with or without IL10 (100 ng/mL) treatment for 12 hours. **(B)** qRT-PCR analysis assessing the expression of *Arg1*, *Mrc1*, and *Socs3* normalized to *GAPDH* in these cells.

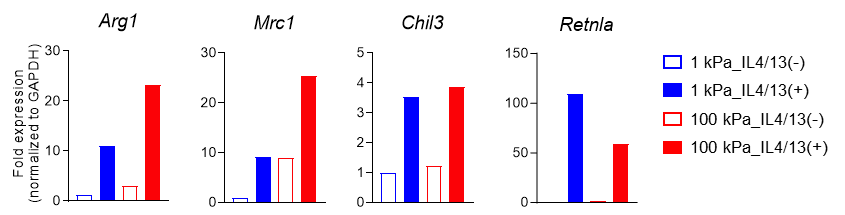

**Supplementary Figure 5. M2 macrophage-associated gene expression is synergistically regulated by cytokines and matrix rigidity.** qRT-PCR analysis for the expression of M2-associated genes (*Arg1*, *Mrc1*, *Chil3*, and *Retnla*) in BMDMs cultured on PA gels of 1 kPa or 100 kPa rigidity, with or without IL4/13 treatment.

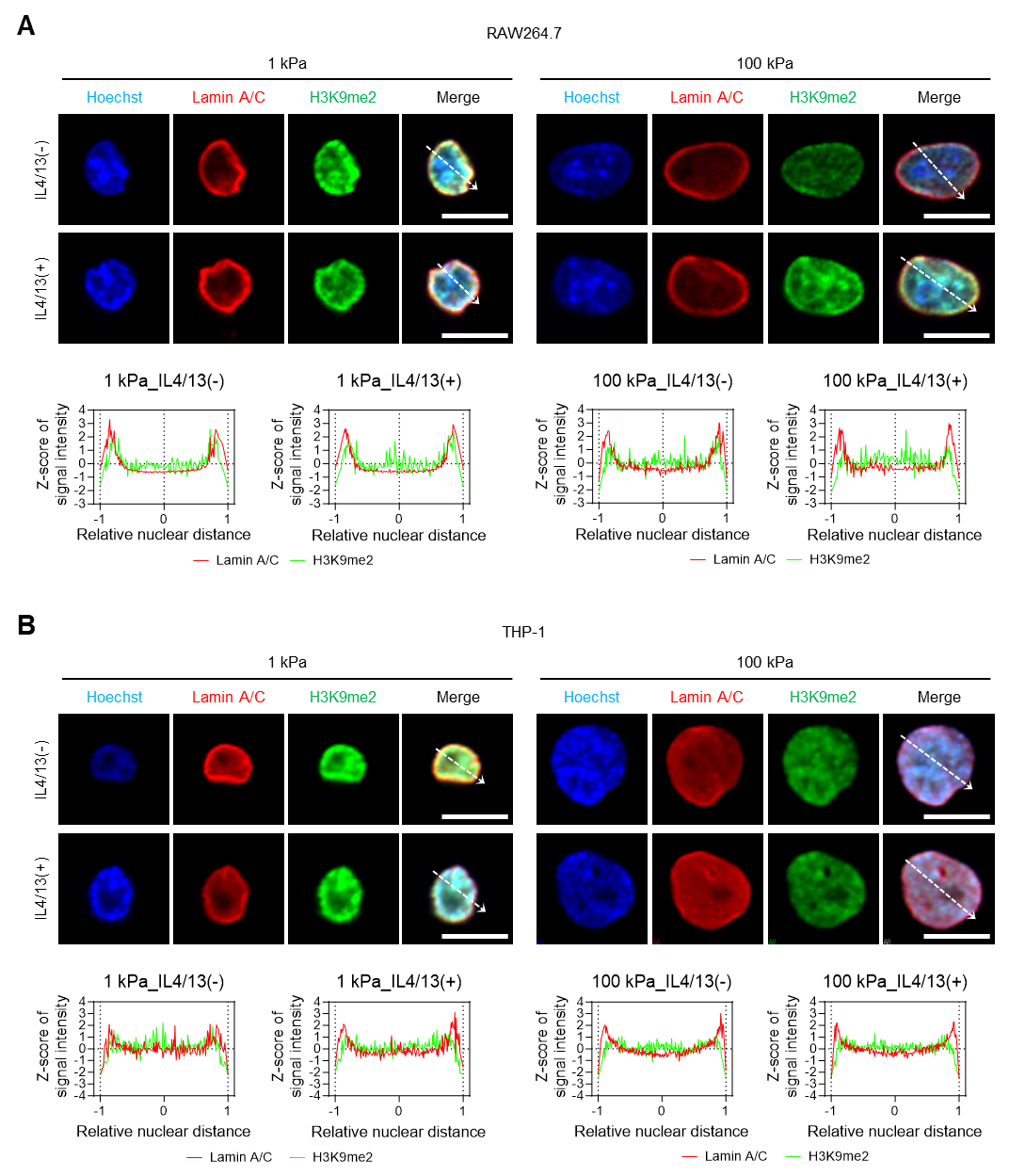

**Supplementary Figure 6. Laminal enrichment of heterochromatin mark, H3K9me2, is attenuated in macrophages by sensing high matrix rigidity.** Representative co-immunostaining images exhibiting Lamin A/C (red), H3K9me2 (green), and Hoechst-33342 (blue) in RAW264.7 and THP-1 cells, cultured on PA gels of either 1 kPa or 100 kPa rigidity, with or without IL4/13 treatment (scale bars = 10 µm). Dashed arrows denote the positions of line signal intensity profiles.

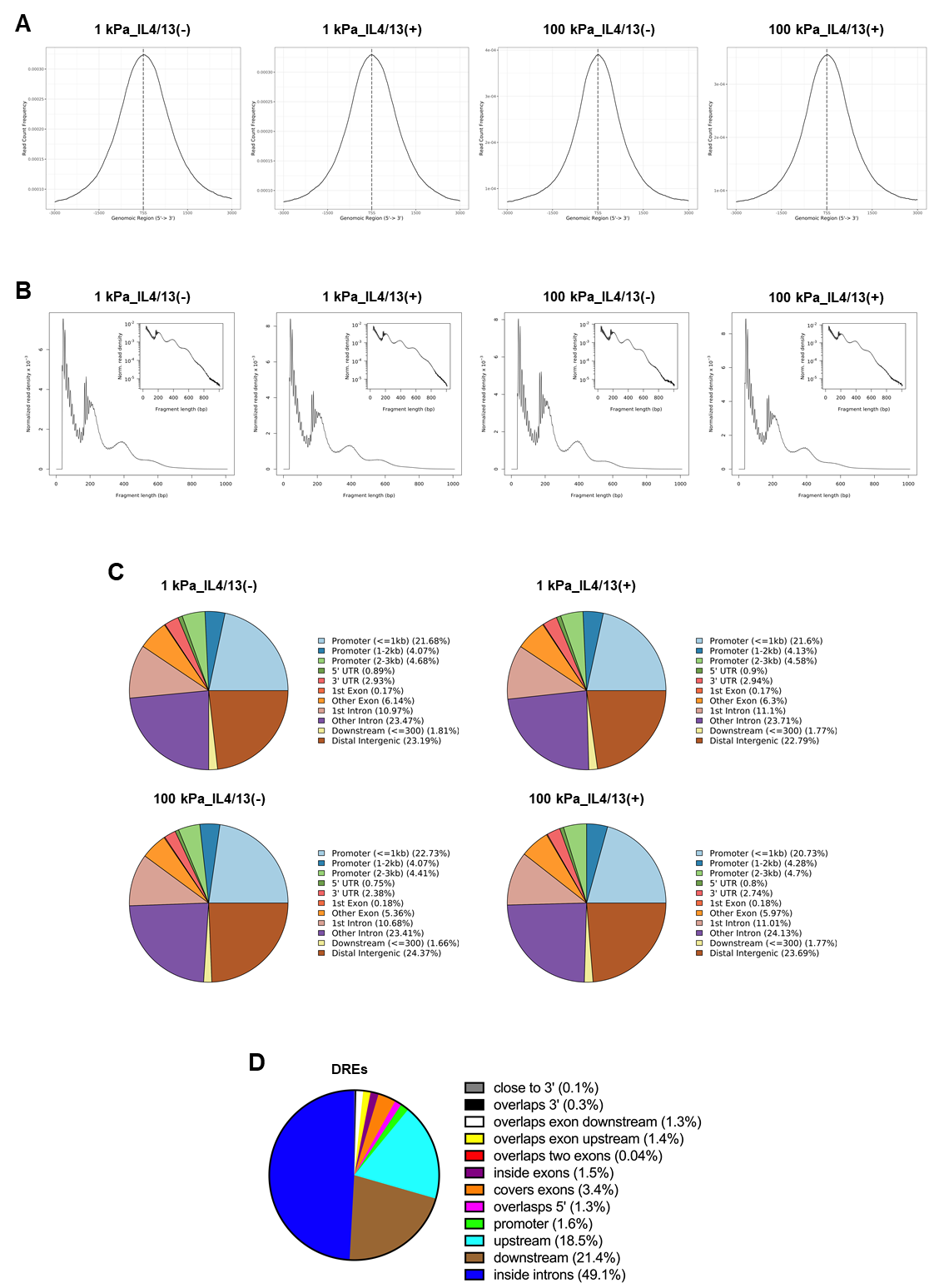

**Supplementary Figure 7. Quality assessment of ATAC-seq dataset.** **(A)** Bell-shaped transcription start site (TSS) enrichment plots showing that read counts are enriched at TSS within genomic regions encompassing ± 3 kbp from TSS in all samples. **(B)** Fragment size distribution plot showing enrichment around 100 and 200 base pair (bp), indicating nucleosome-free and mono-nucleosome-bound fragments. **(C)** Peak annotation pie charts showing that typical patterns of peak distribution in all samples. **(D)** Pie chart showing the peak distribution of distinct regulatory elements (DREs) unique in one or more of the four conditions compared to the other conditions (|fold change|≥1.5, normalized data (log2)≥1).

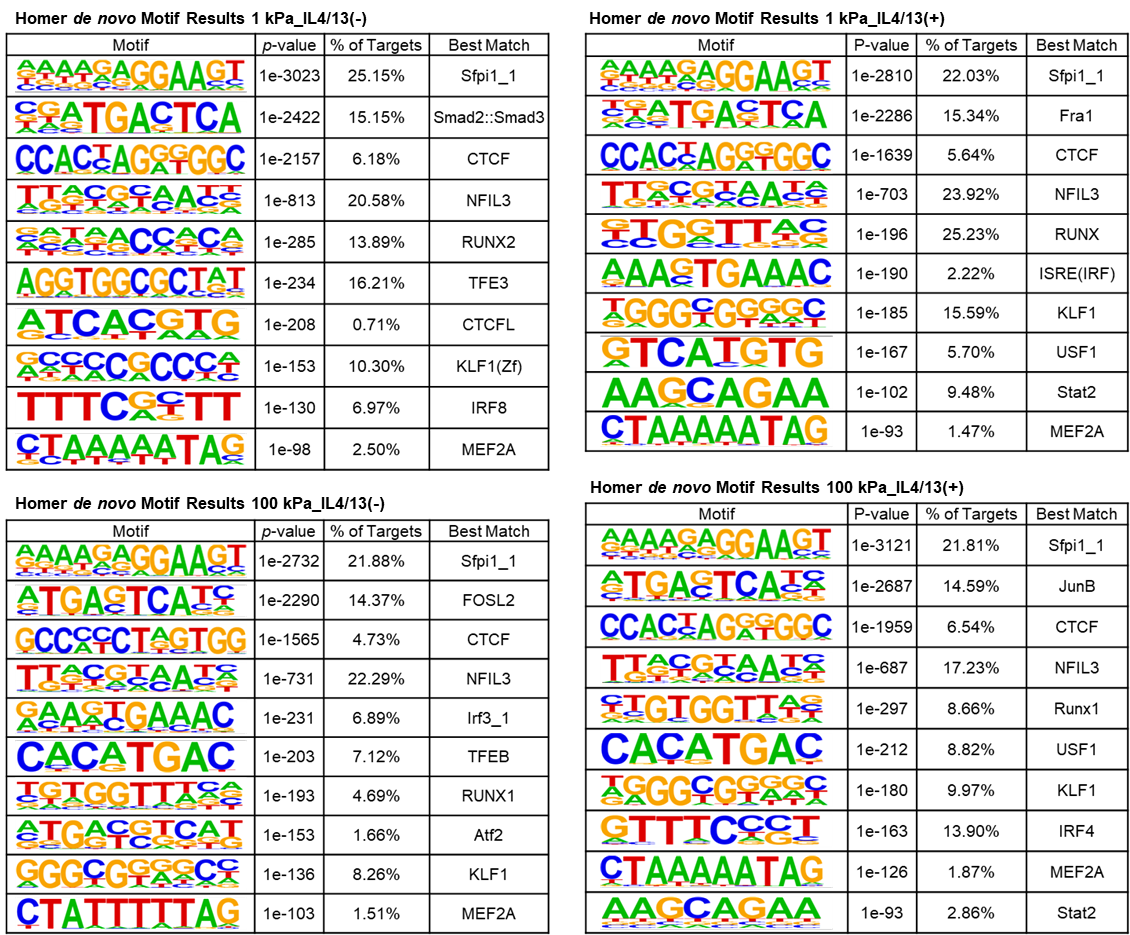

**Supplementary Figure 8. Identification of the top 10 novel transcription factor binding motifs through Homer software across four distinct BMDM culture conditions** (1 kPa_IL4/13(-), 1 kPa_IL4/13(+), 100 kPa_IL4/13(-), and 100 kPa_IL4/13(+)).

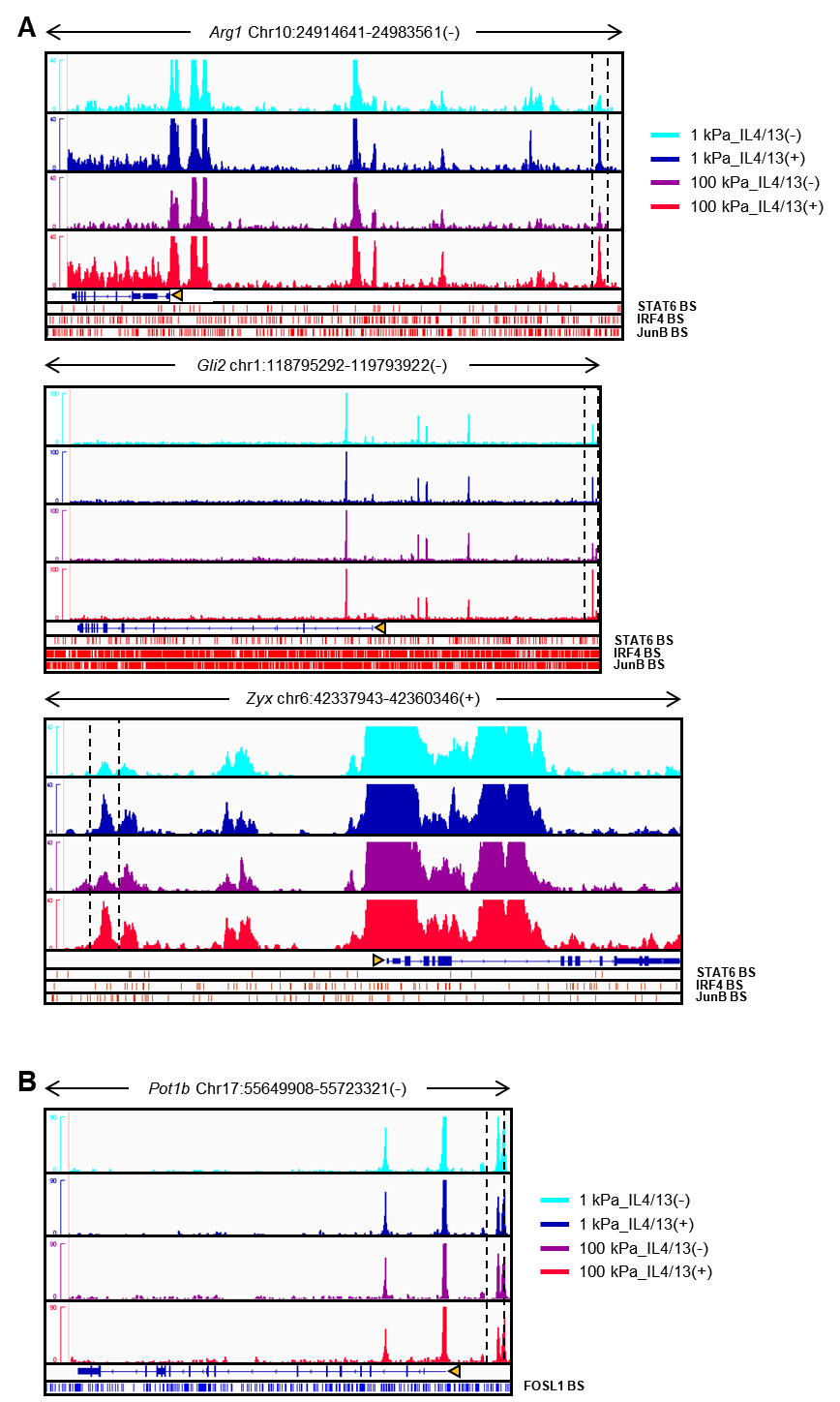

**Supplementary Figure 9. Normalized ATAC-seq profiles across the gene locus in C1_1 and C1_2 clusters.** **(A, B)** Visualization of normalized DRE peaks spanning the chosen gene locus within C1_1 **(A)** and C1_2 **(B)** clusters across the four distinct BMDM culture conditions. Binding sites of M2-activating TFs, including STAT6, IRF4, and JunB, are denoted by red bars, while the binding sites of an M2-inhibiting TF, FOSL1, is represented by blue bars.

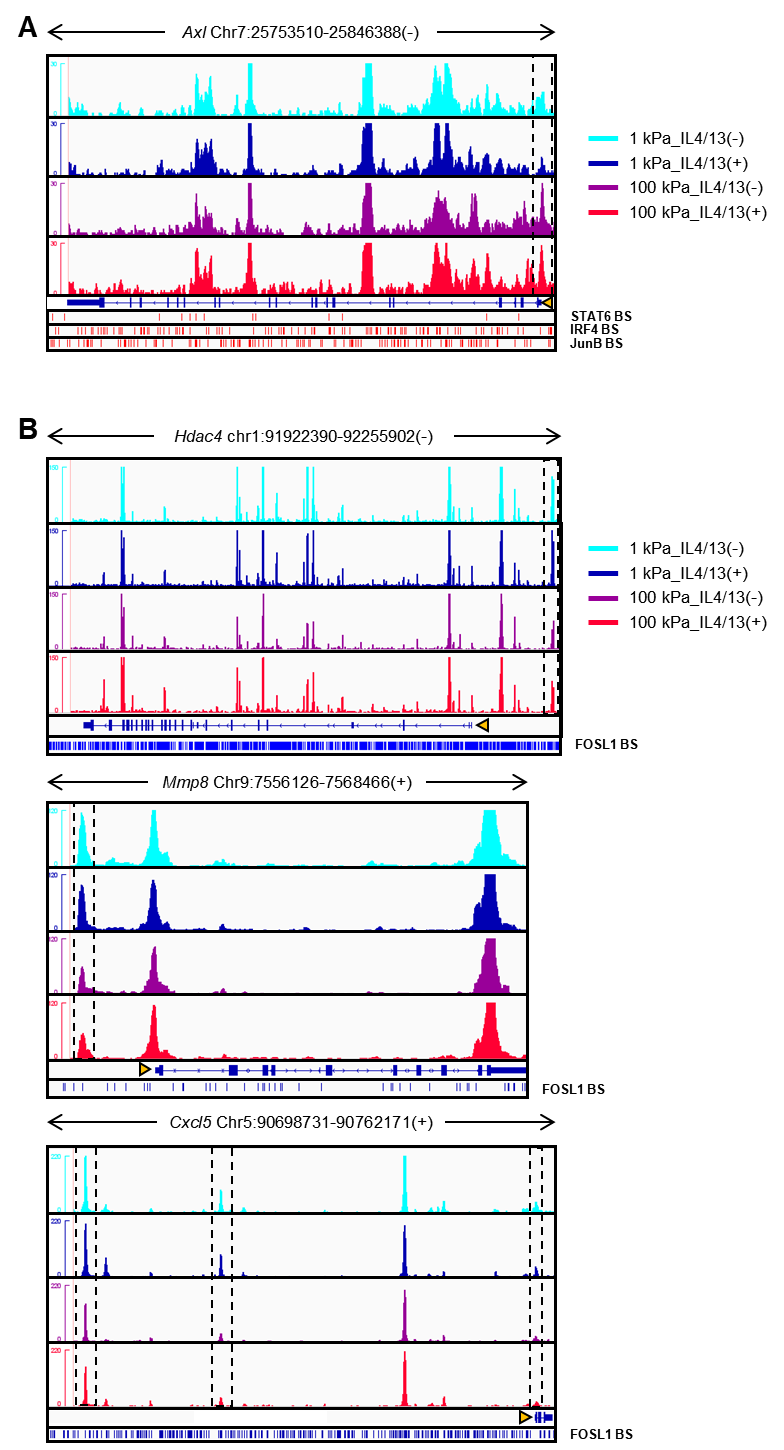

**Supplementary Figure 10. Normalized ATAC-seq profiles across the gene locus in C2_1 and C2_2 clusters.** **(A, B)** Visualization of normalized DRE peaks spanning the chosen gene locus within C2_1 **(A)** and C2_2 **(B)** clusters across the four distinct BMDM culture conditions. Binding sites of M2-activating TFs, including STAT6, IRF4, and JunB, are denoted by red bars, while the binding sites of an M2-inhibiting TF, FOSL1, is represented by blue bars.

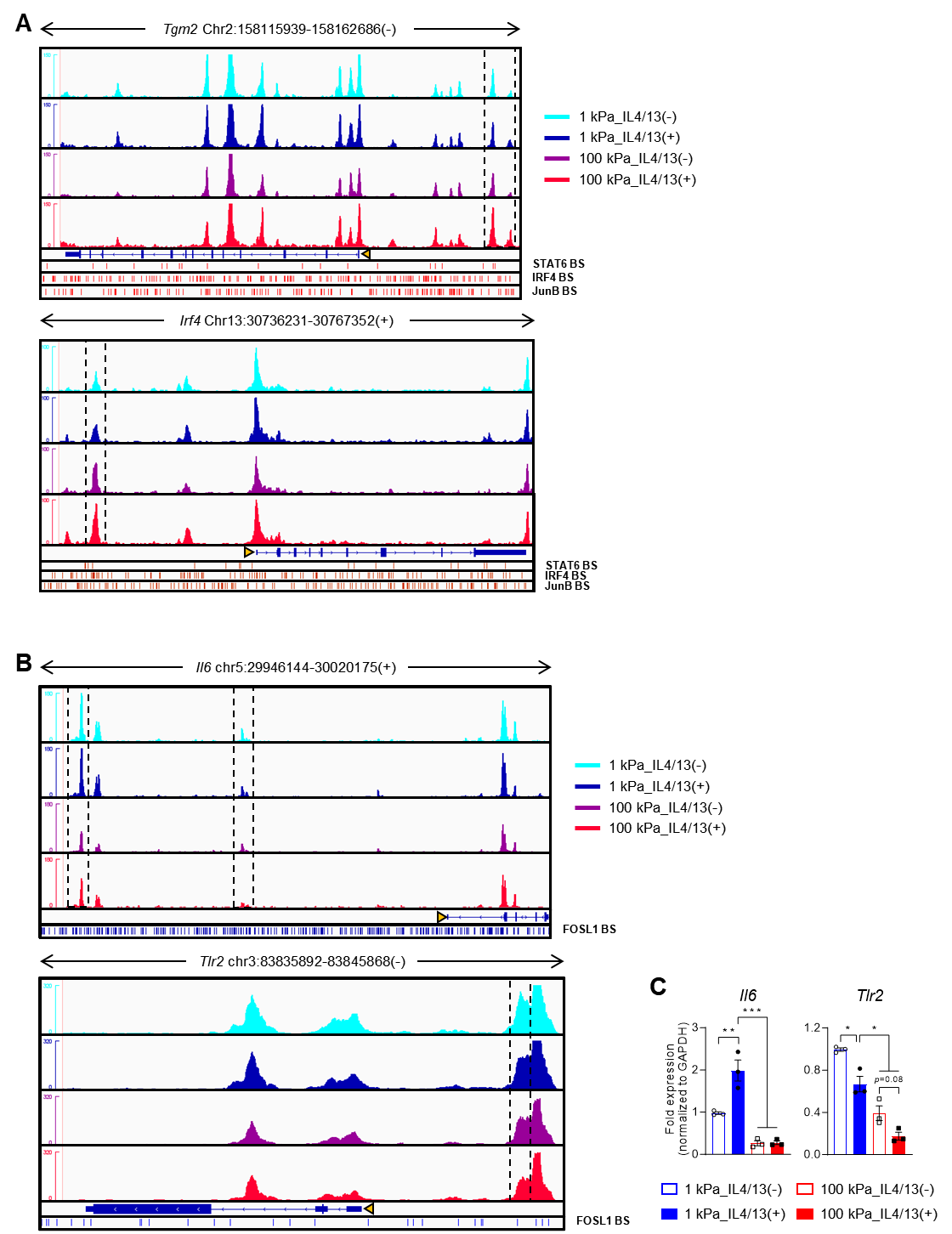

**Supplementary Figure 11. Normalized ATAC-seq profiles across the gene locus in C3_1 and C3_2 clusters.** **(A, B)** Visualization of normalized DRE peaks spanning the chosen gene locus within C3_1 **(A)** and C3_2 **(B)** clusters across the four distinct BMDM culture conditions. Binding sites of M2-activating TFs, including STAT6, IRF4, and JunB, are denoted by red bars, while the binding sites of an M2-inhibiting TF, FOSL1, is represented by blue bars.

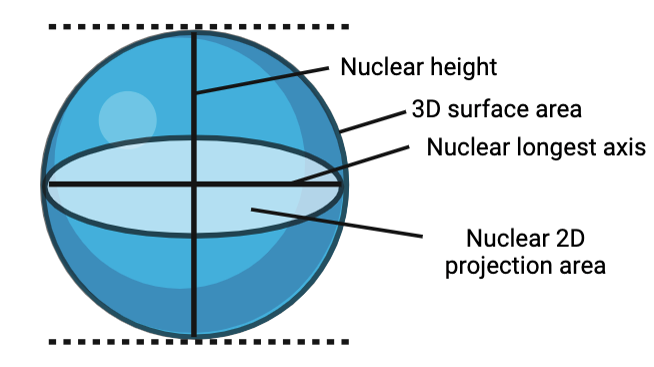

**Supplementary Figure 12.** Nuclear 2D projection area is calculated from a flattened nucleus of plane view. The nuclear flattening index (NFI) is calculated by dividing the length of the longest axis of a nucleus by its height, where a higher NFI value signifies greater nuclear flattening. Nuclear 3D surface area is the total area of nuclear outer surface.

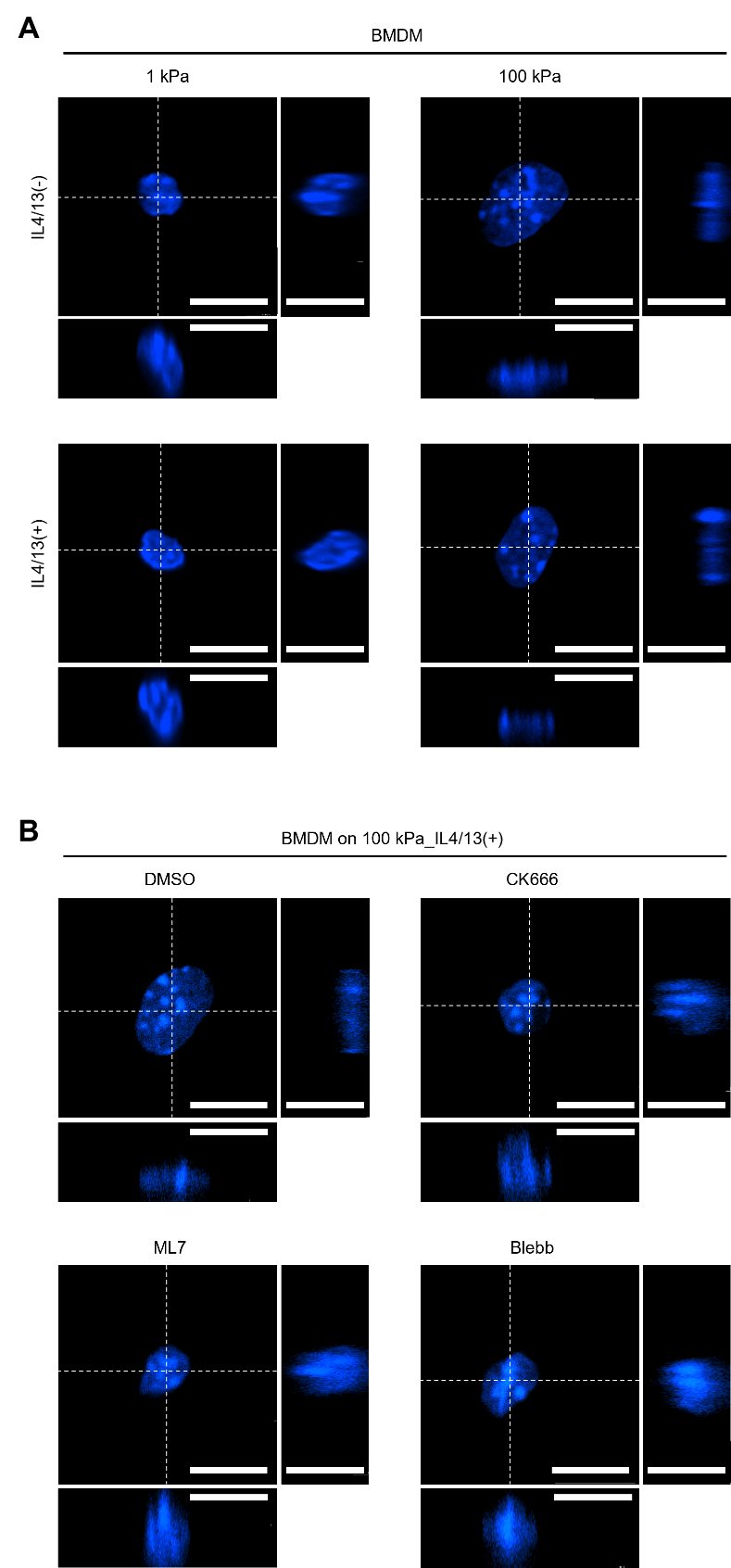

**Supplementary Figure 13. Distinct nuclear morphology of BMDMs on variable substrate rigidity. (A)** Representative top- and side-view images depict the distinct nuclear morphology of BMDMs cultured on either 1 kPa or 100 kPa PA gel in the absence or the presence of IL4/13 (scale bars = 10 µm). **(B)** Representative images of BMDMs cultured on the 100 kPa_IL4/13(+) condition after treatment with CK666, ML7, blebbistatin (Blebb), or DMSO as a vehicle (scale bars = 10 µm). Visualization utilized DAPI (blue) staining. The white dashed lines indicate the sectioning points of side views.

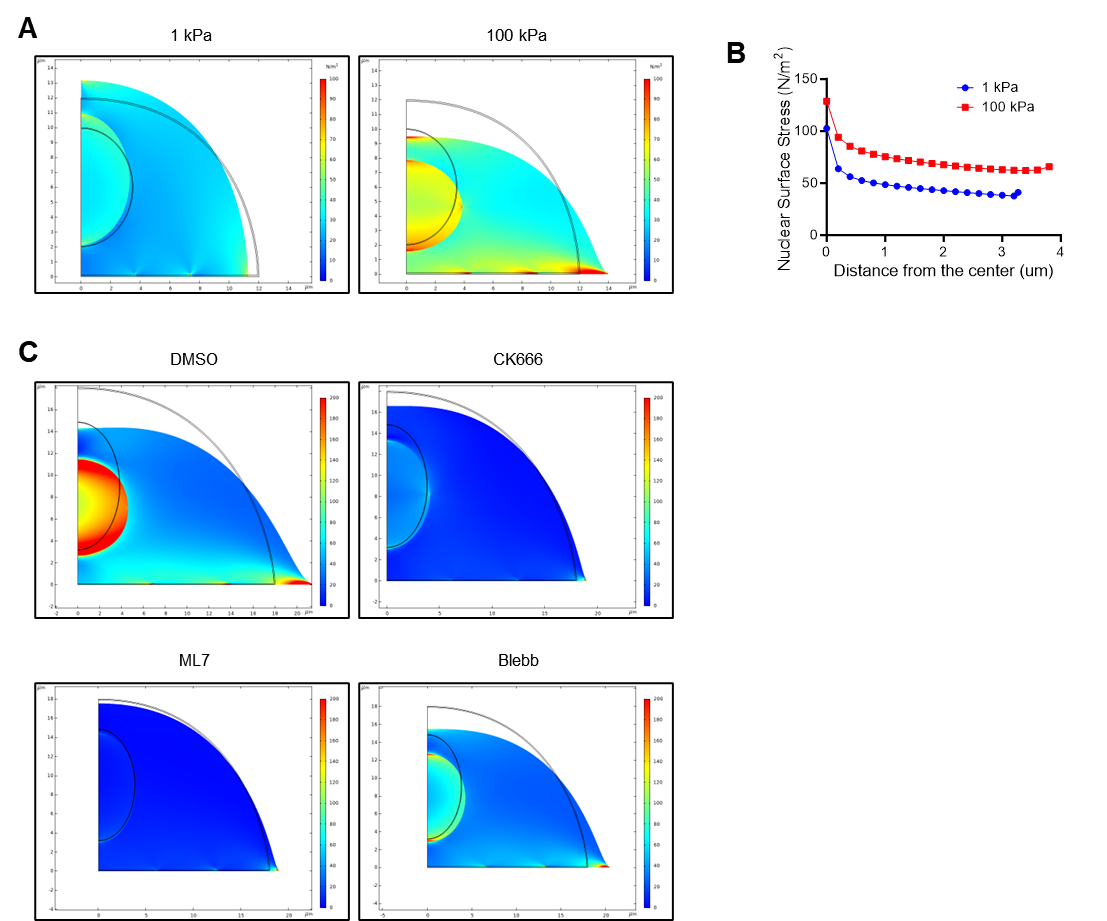

**Supplementary Figure 14. Nuclear deformation and increased nuclear surface stress under high matrix rigidity via actomyosin-mediated force transmission. (A)** 2D sectional views illustrating the deformation and stress distribution within the cell anchored to a compliant (left) or rigid (right) substrate. Cell deformation was simulated through horizontal stretching, and boundary loads were applied to the cytoplasm. The black solid curves indicate the nucleus and cell boundary before deformation, with the nucleus positioned at the center of the hemispherical cell model. Stress distribution after deformation was color-coded. **(B)** Quantification of von-Mises stress on the nuclear surface. von-Mises stresses exhibited a gradual decrease from the center to the boundary of the cell. **(C)** Deformation and stress distribution within the cells anchored to a rigid substrate after treatment with CK666, ML7, Blebbistatin (Blebb), or DMSO as a vehicle.

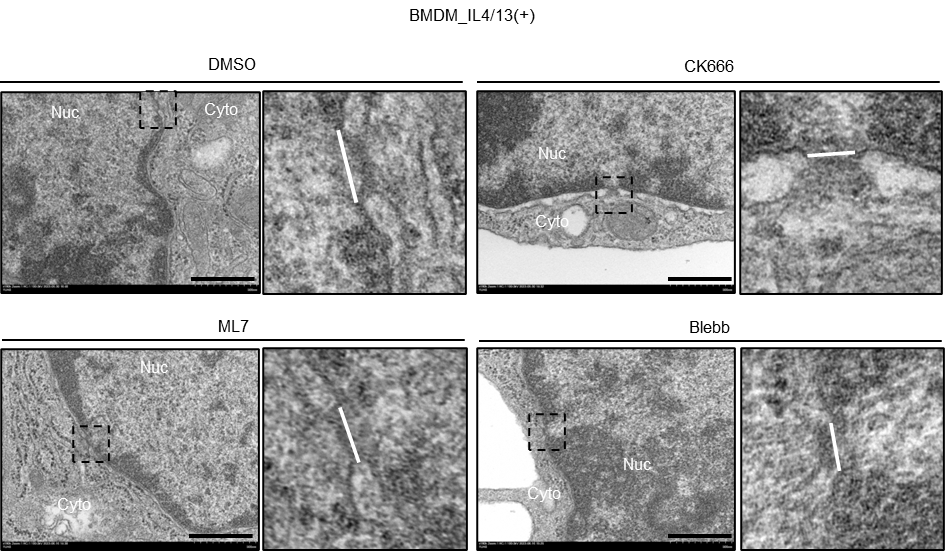

**Supplemental Figure 15. Representative TEM images of nuclear pores in actomyosin-inhibited BMDMs.** Representative TEM images of nuclear pores in BMDMs treated with DMSO, CK666, ML7, or Blebb in the presence of IL4/13 (scale bars = 100 nm). Nuclear pores within dashed boxes are captured at a higher magnification. The lengths of open nuclear pores are indicated by white lines. Nuc, nucleus; Cyto, cytoplasm.

**Supplementary Table 1. Mechanical properties of a model cell.**

| Property | Cytosol | Nucleus | Unit | Reference |
| --- | --- | --- | --- | --- |
| Shear modulus | 50 | 80 | Pa | ^1-3^ |
| Poisson’s ratio | 0.4 | 0.4 | - | ^2-5^ |
| Density | 1 | 1.2 | kg/m^3^ | ^6,7^ |
| Young’s modulus | 200 | 300 | Pa | ^8^ |

**Supplementary Table 2. Primer sequences used in qRT-PCR.**

| Mouse gene | Sequence |
| --- | --- |
| *Arg1* | Forward: 5’-TTATCGGAGCGCCTTTCTCA-3’ |
|  | Reverse: 5’-AGACCGTGGGTTCTTCACAA-3’ |
| *Mrc1* | Forward: 5’-TGCCAAGTGGGAAAATCTGG-3’ |
|  | Reverse: 5’-TAGGGCCACCACTGATTAGG-3’ |
| *Chil3* | Forward: 5’-GAAGCTCTCCAGAAGCAATCCT-3’ |
|  | Reverse: 5’-CTGGTAGGAAGATCCCAGCTGTA -3’ |
| *Retnla* | Forward: 5’-CCAATCCAGCTAACTATCCCTCC-3’ |
|  | Reverse: 5’-CCAGTCAACGAGTAAGCACAG-3’ |
| *Il10* | Forward: 5’-GTGGAGCAGGTGAAGAGTGAT-3’ |
|  | Reverse: 5’-AGTCCAGCAGACTCAATACACA-3’ |
| *Tnfα* | Forward: 5’-CCACGCTCTTCTGTCTACTG-3’ |
|  | Reverse: 5’-CTGATGAGAGGGAGGCCATT-3’ |
| *Il6* | Forward: 5’-CTTCACAAGTCGGAGGCTTAAT-3’ |
|  | Reverse: 5’-ACTCCAGGTAGCTATGGTACTC-3’ |
| *Jak2* | Forward: 5’-TTGTGGTATTACGCCTGTGTATC-3’ |
|  | Reverse: 5’-ATGCCTGGTTGACTCGTCTAT-3’ |
| *Tgfbi* | Forward: 5’-CATTGGCACCAACAAGAAATAC-3’ |
|  | Reverse: 5’-CTTTTCATATCCAGGACAGCAC-3’ |
| *Pparg* | Forward: 5’-GGCCTCCCTGATGAATAAA-3’ |
|  | Reverse: 5’-GCTCCATAAAGTCACCAAAG-3’ |
| *Socs3* | Forward: 5’-CCTTTGACAAGCGGACTCTC-3’ |
|  | Reverse: 5’-GCCAGCATAAAAACCCTTCA-3’ |
| *Cxcl5* | Forward: 5’-GGTCCACAGTGCCCTACG-3’ |
|  | Reverse: 5’-GCGAGTGCATTCCGCTTA-3’ |
| *Tlr2* | Forward: 5’-ACTTCTCTGCTTTTCGTTCATC-3’ |
|  | Reverse: 5’-CTCGTAGCATCCTCTGAGATTT-3’ |
| *GAPDH* | Forward: 5’-CTGCACCACCAACTGCTTAG-3’ |
|  | Reverse: 5’-GTCTTCTGGGTGGCAGTGAT-3’ |

**Supplementary Table 3. Antibodies used in Western blot.**

| Target antigen | Vendors or Source | Catalog number | Dilution information |
| --- | --- | --- | --- |
| Phospho-STAT6 | Cell Signaling | 56554S | 1:1000 |
| STAT6 | Cell Signaling | 5397S | 1:1000 |
| Phospho-STAT3 | Cell Signaling | 9145S | 1:1000 |
| STAT3 | Cell Signaling | 4904S | 1:1000 |
| ARG1 | Invitrogen | PA5-29645 | 1:1000 |
| H3K9me2/3 | Cell Signaling | 5327S | 1:500 |
| B-tubulin | Cell Signaling | 2146S | 1:1000 |
| B-actin conjugated with peroxidase | Cell Signaling | A3854 | 1:10000 |

**Supplementary Table 4. Antibodies used in immunofluorescent staining.**

| Target antigen | Vendors or Source | Catalog number | Dilution information |
| --- | --- | --- | --- |
| Phospho-STAT6 | Cell Signaling | 9361S | 1:400 |
| Lamin A/C | Cell Signaling | 4777S | 1:200 |
| Lamin B1 | Abcam | Ab16048 | 1:200 |
| VINCULIN | Abcam | ab129002 | 1:1000 |
| H3K9me2/3 | Cell Signaling | 5327S | 1:200 |
| H3K9me2 | Invitrogen | PA5-16195 | 1:200 |
